# ATP driven diffusiophoresis: active cargo transport without motor proteins

**DOI:** 10.1101/2020.05.01.072744

**Authors:** Beatrice Ramm, Andriy Goychuk, Alena Khmelinskaia, Philipp Blumhardt, Kristina A. Ganzinger, Erwin Frey, Petra Schwille

## Abstract

Morphogenesis and homeostasis of biological systems are intricately linked to gradient formation through energy dissipation. Such spatial organization may be achieved via reaction-diffusion or directional cargo transport, as prominently executed by motor proteins. In contrast to these processes that rely on specific protein interactions, active transport based on a non-specific, purely physical mechanism remains poorly explored. Here, by a joint experimental and theoretical approach, we describe a hidden function of the MinDE protein system from *E. coli:* Besides forming dynamic patterns, this system accomplishes the active transport of large, functionally unrelated cargo on membranes *in vitro*. Remarkably, this mechanism allows to sort diffusive objects according to their effective size, as evidenced using modular DNA origami–streptavidin nanostructures. We show that the diffusive fluxes of MinDE and cargo couple via density-dependent friction. This non-specific process constitutes a Maxwell-Stefan diffusiophoresis, so far undescribed in a biologically relevant setting. Such nonlinear coupling between diffusive fluxes could represent a generic physical mechanism for the intracellular organization of biomolecules.

## Introduction

Spatiotemporal organization is a hallmark of living cells. It generally emerges through redistribution and transport of cargo molecules via motor proteins^1^, self-assembling cytoskeletal elements^2^ or self-organizing reaction-diffusion systems^3^. Coupling of cargo to the energy-dissipating NTPases that drive the transport is usually mediated by specific protein-protein interactions. Non-specific coupling of biomolecules, on the contrary, is poorly explored in biology and so far only a few examples of molecular transport based on purely physical mechanisms have been reported: For instance, a study in mouse oocytes showed that active diffusion of actin-coated vesicles generated a pressure gradient that positioned large objects like the nucleoid in the cell centre^4,5^. In the *Caenorhabditis elegans* zygote cortical flows were shown to couple to the PAR reaction-diffusion system via advective transport^6^. Another recent example comes from *in vitro* studies of the *Escherichia coli* Min system: MinDE self-organization induced patterns and gradients of inert membrane-bound molecules^7,8^.

The Min system regulates the site of cell division in *E. coli* and has become a paradigmatic model for pattern formation in biology^9–11^. Despite its compositional simplicity, the system exhibits rich dynamics that have been explored *in vivo*^9,11,12^, reconstituted *in vitro*^13–15^ and described by physical theories^16–20^. The core of this reaction-diffusion system consists of only two proteins, the ATPase MinD and the ATPase activating protein MinE, which interact and reversibly bind to the membrane^11,13^. In the rod-shaped *E. coli*, MinDE oscillate from pole to pole^10–12^. *In vitro*, these two proteins form travelling surface waves and quasi-stationary patterns on planar artificial membranes^13–15^ or also exhibit oscillations when geometrically confined^21,22^. MinDE dynamics can provide spatial cues for particular proteins: MinC specifically binds to MinD and thus follows its movements^10,21,23–25^. In turn, MinC constrains the localization of the main divisome protein FtsZ, by inhibiting its polymerization^26,27^.

Besides this well-described patterning by specific interactions with clear physiological evidence, MinDE self-organization has recently shown an intriguing hidden function *in vitro:* MinDE regulated the localization of unrelated membrane-bound molecules (cargo) in space and time in the absence of MinC/FtsZ^7,8^. The ability of MinDE to redistribute arbitrary molecules suggested that MinDE could further enhance cell division by prepositioning membrane proteins to the cell middle. Even more importantly, it also hinted towards a generic active transport mechanism that neither requires motor proteins nor specific protein interactions. Nevertheless, the underlying physics and the broader biological implications remained unknown.

Here, we set out to decipher the physical mechanism underlying this MinDE-dependent transport phenomenon through a joint experimental and theoretical investigation. We quantitatively probed MinDE-dependent transport with a synthetic cargo based on membrane-anchored composite nanostructures consisting of a DNA origami scaffold and streptavidin building blocks. By varying the number of streptavidin, we found that MinDE-induced transport depends on the effective size of the cargo and leads to spatial sorting of different cargo molecules. Theoretical analysis of these data demonstrated that cargo particles are carried along the diffusive fluxes of MinD proteins via an effective density-dependent inter-particle friction, which we term Maxwell-Stefan diffusiophoresis. In general, diffusiophoresis refers to particle transport by concentration gradients of small solutes, which has been predicted and demonstrated for three-component gases^28–31^ and colloidal suspensions^32–37^. In a biologically relevant context, however, so far only theoretical studies have suggested that gradients of small molecules, such as ATP, could induce metabolism-dependent transport of large particles^38^. We show in both experiment and theory that Maxwell-Stefan diffusiophoresis, powered by the inhomogeneous distribution of energy-dissipating NTPases, can drive the redistribution of similar-sized as well as much larger biomolecules. Importantly, Maxwell-Stefan diffusiophoresis potentially represents a generic active transport mechanism in cells. Since prokaryotes lack specialized motor proteins, such a mechanism could be particularly important for this branch of life, and it might have been prevalent in early stages of life on earth.

## Probing the transport mechanism with a synthetic cargo

Conceptually, we asked whether cargo transport by MinDE arises from thermodynamic forces or requires active processes. To experimentally address this question and test possible mechanisms, we set up a highly controllable *in vitro* platform. We reconstituted MinDE pattern formation on supported lipid bilayers (SLB), as previously established^13^. Here, we chose conditions under which MinDE form quasi-stationary labyrinth patterns to simplify the assay^15^. Although these MinDE patterns are in a quasi-steady state, they are maintained by the same nonequilibrium dynamics as propagating MinDE surface waves: MinE induces MinD membrane detachment and so there are persistent cyclic protein fluxes between bulk solution and membrane^15,24^. To be able to quantitatively assess the interaction of MinDE with cargo molecules on the membrane, we employed a synthetic cargo. This cargo consisted of a DNA origami nanostructure as scaffold and streptavidin molecules that serve as modular building blocks and connectors to the membrane (Fig. 1a). In particular, the origami (20 helix bundle; 110×16×8 nm)^39^ featured 7 dyes on the upper facet for visualization and 42 modifiable sites at the bottom facet. The bottom positions could be specifically addressed for the incorporation of biotinylated oligonucleotide handles. These handles in turn bound to streptavidin coupled to biotinylated lipids in the SLB (Fig. 1a).

**Fig. 1:**
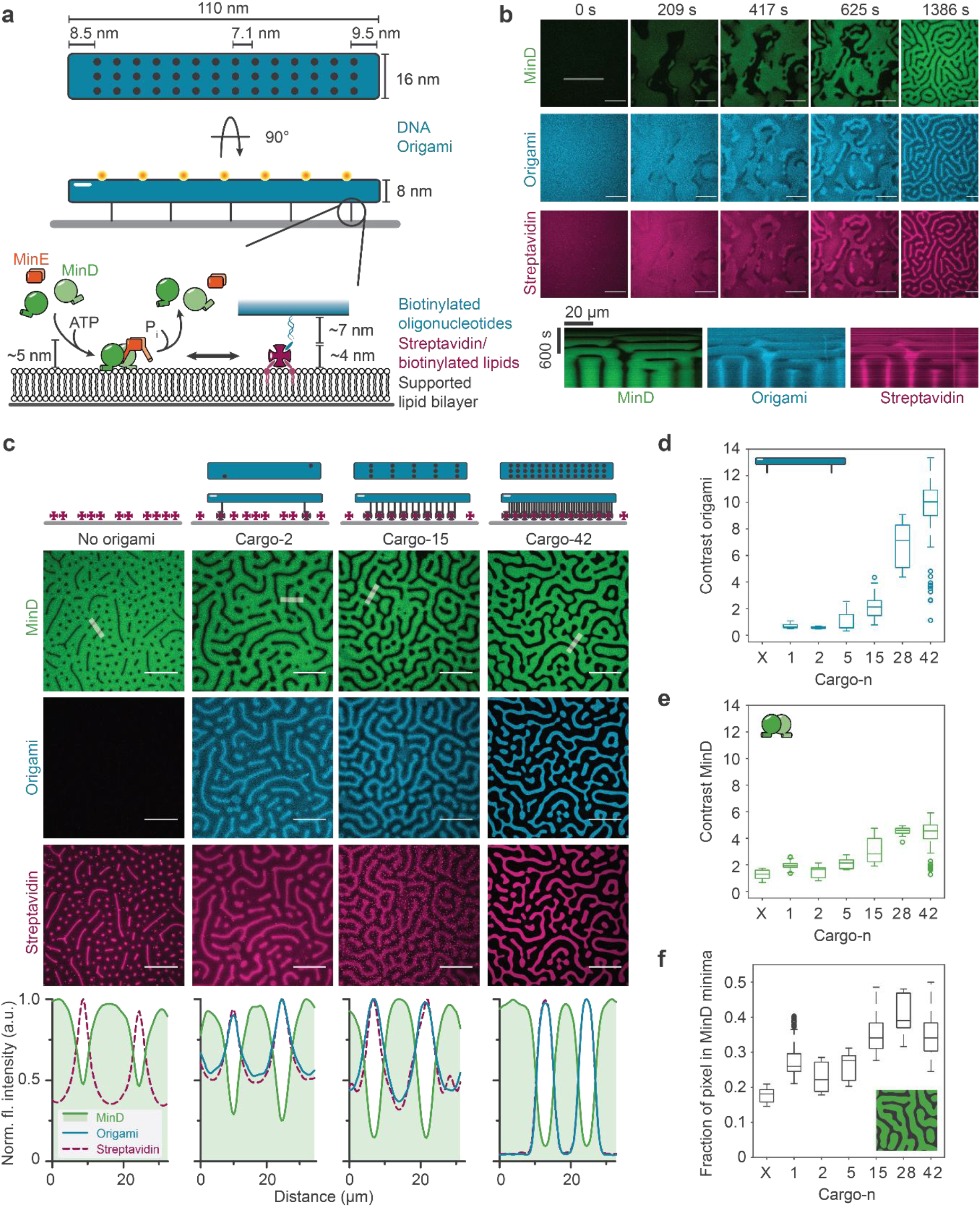
MinDE-driven cargo unmixing depends on the effective size of the cargo. **a**, Schematic of the synthetic, membrane-anchored cargo consisting of a DNA origami scaffold and streptavidin building blocks. DNA origami nanostructure illustrating the position of 7 dyes at the upper facet and 42 addressable sites for incorporation of biotinylated oligonucleotides at the lower facet. Biotinylated oligonucleotides are then bound to lipid-anchored streptavidin on the SLB. **b**, Representative time-series and kymograph (along line selection) of MinDE self-organization inducing patterns of cargo-2, i.e. of DNA origami with two bound streptavidin (1 μM MinD (30% EGFP-MinD), 1.5 μM MinE-His, 0.1 nM origami-Cy5 with 2 biotinylated oligonucleotides, Alexa568-streptavidin). **c**, Representative images and fluorescence intensity line plots (smoothed) of established MinDE labyrinth patterns and anti-correlated DNA origami and streptavidin patterns when no origami, cargo-2, cargo-15 or cargo-42 is present. Contrast of the resulting patterns and size of the MinD minima increases with increasing number of streptavidin. Box plot of the contrast of **d**, cargo and **e**, MinD patterns, and **f**, of the fraction of pixel classified as MinD minima, when no origami, cargo-2, cargo-15 or cargo-42 is present. Box plot lines are median, box limits are quartiles 1 and 3, whiskers are 1.5× interquartile range (IQR) and points are outliers. Data from at least two independent experiments with in total number of analyzed images per condition N(No origami)=32, N(Cargo-1)=112, N(Cargo-2)=41, N(Cargo-5)=48, N(Cargo-15)=110, N(Cargo-28)=32, N(Cargo-42)=98. Scale bars: 50 μm

When MinDE self-organization was started with ATP in the presence of this synthetic cargo, the cargo components, origami structures and streptavidin, reorganized into patterns from an initially homogenous state (Fig. 1b, Fig. S1, Supplementary Movie 1). They always gathered in MinD-depleted regions, forming patterns that were superimposable and anti-correlated to the MinDE patterns. As expected DNA origami and streptavidin co-localized, hence, in the following analyses origami fluorescence serves as a proxy for cargo localization. Cargo accumulation in MinD-depleted regions was evident throughout the entire duration of the experiment, from initiation of self-organization to the establishment of the final pattern (Fig. 1b). Similarly, when we altered the established MinDE/cargo patterns by adding more MinE, the cargo channel reflected the changes in MinDE patterns, moving in an anti-correlated fashion (Fig. S2, Supplementary Movie 2). In contrast, when MinE was omitted from experiments, MinD and the cargo molecules remained uniformly distributed (Fig. S3). These findings indicate that the spatial heterogeneity of cargo distribution is not caused by depletion forces, such as in filament bundling^40^: depletion forces should lead to aggregation of large particles (cargo) even in a homogeneous field of smaller particles (MinD)^41^. Furthermore, depletion forces would imply a biased motion of cargo towards regions of high MinD density, where they would preferably agglomerate, which we also do not observe (see SI for details). Hence, our data show that cargo transport requires the presence of MinD, MinE and ATP, and thus active MinDE self-organization and pattern formation.

## Effective cargo size determines the extent of cargo unmixing

Having shown that MinDE are capable of redistributing our synthetic cargo, we next set out to systematically vary the interaction of the cargo molecules with MinDE. To that end, we took advantage of the high addressability of the DNA origami scaffold and the composite nature of our cargo. MinD as well as the lipid-anchored streptavidin form a monomolecular layer of about 5 nm height on the membrane^42,43^, whereas the origami scaffolds are located at an altitude of about 5 and 11 nm above the membrane (see Supplementary Note 1). Therefore, MinDE move on the membrane below the altitude of the origami scaffold and should mainly interact with the membrane-bound streptavidin. Hence, varying the number of streptavidin bound to the origami scaffold allowed us to achieve fine control and a large dynamic range of effective cargo size, and thus its interaction with MinDE, but also its diffusion on the membrane. In the following we refer to these composite objects, i.e. DNA-origami with *n* ∈ {1,2,5,15,28,42} streptavidin, as “cargo-n” (Fig. 1).

To quantitatively assess the interaction between MinDE and the respective cargo, we analyzed the final quasi-stationary patterns of cargo and MinD (Fig. 1c, Fig. S4). Specifically, as a measure for the enrichment of the respective molecules, we determined the fluorescence intensity contrast, (*I*_max_ − *I*_min_)/*I*_max_. We found that the cargo patterns became much sharper, i.e. exhibited larger contrast, with increasing cargo size (Fig. 1d, Fig. S5). This increase in contrast of cargo patterns was accompanied by sharper and also narrower MinDE patterns, as indicated by an increased region of pixels classified as MinD minima (Fig. 1e,f). Thus, MinDE dynamics dictate the localization of cargo on the membrane in a size-dependent manner, but are in turn also impacted by their presence. Cargo distribution in a static gradient of accessible space should not depend on the properties of cargo (see SI for details). Therefore, even though the diffusion of MinD on the membrane under the given conditions is very slow (*D* = 0.013 μ*m*^2^ *s*^−1^)^24^, we conclude that MinD proteins do not act as static obstacles on the membrane that would bias cargo diffusion via static volume exclusion, a second option for a thermodynamic force.

## Thermodynamic forces cannot explain MinDE-dependent cargo transport

As our experimental data disqualified both depletion forces and static volume exclusion as possible explanations for cargo redistribution, we wondered whether mobile MinD proteins could transport cargo via entropic mixing effects, a third option for a thermodynamic force. To test this hypothesis, we formulated a quantitative Flory-Huggins-type model (without any fitting parameters; see Table S1 for particle densities) taking into account the following considerations: (i) Each origami scaffold crosslinks *n* streptavidin blocks into a passive polymer-like object on the membrane (cargo-n); (ii) Unless the number of cargo molecules on the membrane is limited by the abundance of membrane-bound streptavidin (e.g. for *n* > 15; see SI and Fig. S5), the remainder of streptavidin molecules is free and hence moves independently of the cargo molecules. With these constraints we asked: given a heterogeneous distribution of active particles (MinD proteins), what is the equilibrium distribution of passive particles (cargo and free streptavidin molecules) in between them? To answer this question, we calculated the corresponding chemical potentials *μ*_*i*_ for each species (see SI). Furthermore, we assumed that the passive particles, cargo *c*_*c*_ and free streptavidin *c*_*s*_, adopt a thermal equilibrium state with vanishing gradients in the chemical potentials (**∇***μ*_*c*_ = **∇***μ*_*s*_ = 0) in an adiabatic response to the imposed fixed distribution of active particles (MinD proteins *c*_*p*_; **∇***μ*_*p*_ ≠ 0). Our theoretical analysis shows that entropic mixing effects can in principle lead to transport of passive particles in a gradient of active particles (Fig. 2b,c). However, experimentally we observed a far stronger redistribution of the passive cargo molecules than entropic mixing would predict (Figure 2b). In other words, the weak entropic sorting of the cargo’s small streptavidin blocks in a gradient of MinD proteins was not sufficient to overcome the strong entropic repulsion between the large origami scaffolds. Consequently, we also rejected entropic mixing in fixed external (chemical potential) gradients as the mechanism underlying the MinDE-dependent cargo transport.

**Fig. 2:**
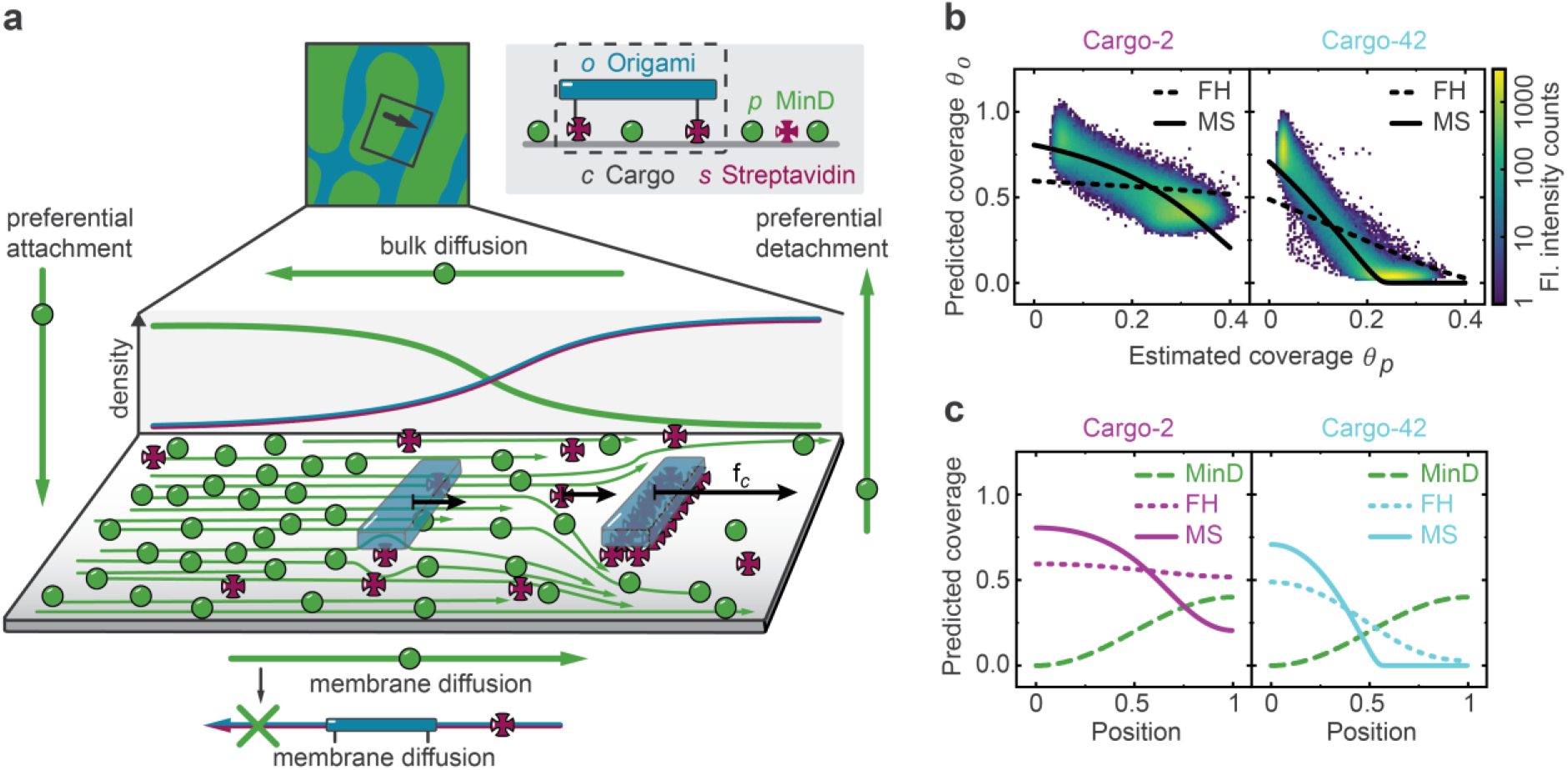
MinDE-dependent cargo transport is explained by friction between particle fluxes, not by mixing or size effects. **A**, Schematic of our theoretical model. Reactions and diffusion of MinDE proteins generate MinD (green) density gradients. Far away from the membrane, the gradients must point in the opposite direction as on the membrane. MinD protein gradients induce diffusive MinD fluxes (green arrows) on the membrane, which exert a frictional force (black arrow) on the cargo molecules (blue origami scaffold with magenta streptavidin building blocks). As a result, a cargo molecule gradient builds up. In steady state, thermodynamic forces (entropic repulsion and mixing, which drive cargo diffusion) balance the frictional force that is exerted by protein flows. **b**, Cross-correlation function between MinD coverage (*θ*_*p*_) and origami coverage (*θ*_*c*_), for two different cargo species, cargo-2 and cargo-42. The color-coded 2D-histogram represents our experimental data, while the solid and dashed lines correspond to two candidate models. The Flory-Huggins type model (FH), whose parameters are fully determined by our experiments, fails to account for cargo transport: the weak entropic sorting of streptavidin blocks in an external gradient of proteins is not sufficient to overcome the strong repulsion of the bulky DNA origami scaffolds. Instead, we find that the Maxwell-Stefan type model (MS), with an estimated interaction parameter, explains cargo transport. **c**, Spatial distribution of cargo molecules in response to the (imposed) MinD profile, corresponding to the cross-correlation functions in b. The Maxwell-Stefan type model allows for stronger reorganization of cargo than the Flory-Huggins type model: in addition to thermodynamic forces, cargo transport is further driven by frictional coupling to MinD protein fluxes. Model parameters: (cargo-2) average coverages 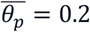, 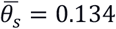 and 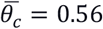; interaction parameter (in terms of MinD coverage) 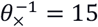; (cargo-42) average coverages 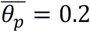, 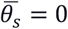 and 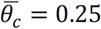; interaction parameter (in terms of MinD coverage) 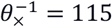. Surface coverages *θ* = *a c* and surface densities *c* are connected by the particle size *a*.

## Friction between particle fluxes accounts for cargo transport

Taken together, we find that none of the in principle feasible thermodynamic mechanisms can actually explain MinDE-induced cargo transport, which suggested that the underlying mechanism is genuinely nonequilibrium in nature. Therefore, we relaxed our previous assumption of fixed external chemical potential gradients and considered their dynamics. According to Onsager’s theory of nonequilibrium thermodynamics^44^, gradients in a chemical potential **∇***μ*_*i*_ imply particle fluxes ***j***_*i*_. In the present context, a possible candidate for a non-equilibrium process in a crowded environment is the coupling of particle fluxes through friction caused by non-specific interactions between proteins and cargo molecules on the membrane. Such coupling between diffusive fluxes has been predicted by the Maxwell-Stefan theory of diffusion^28,29^ and has been experimentally observed for three-component gas mixtures^30,31^. The theory asserts that each species on the membrane obeys an effective force-balance equation^28,29^:

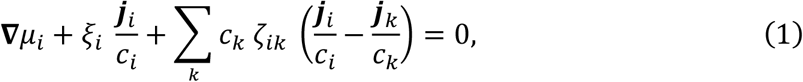

where the index *i* ∈ {*p, c, s*} refers to MinD proteins, cargo molecules with bound streptavidin, and free streptavidin molecules, respectively. In the nonequilibrium steady state, thermodynamic driving forces, caused by chemical potential gradients, are balanced by friction forces between the different macromolecules and lipid molecules (with friction coefficient *ξ*_*i*_) as well as among the macromolecules themselves (with coupling constants *ζ*_*ik*_). While cargo and streptavidin molecules exhibit Brownian motion and relax to a thermal equilibrium state with vanishing fluxes ***j***_*c*_ = ***j***_*a*_ = 0, the MinD protein patterns are kept in a non-equilibrium steady state maintained by off-equilibrium chemical reactions (ATPase activity). In the absence of mutual friction between the macromolecules (*ζ*_*ik*_ = 0), Eq. 1 reduces to the Flory-Huggins model (**∇***μ*_*c*_ = **∇***μ*_*s*_ = 0) which implies weak cargo redistribution in a static gradient of active proteins. In contrast, in the presence of (frictional) coupling (*ζ*_*ik*_ ≠ 0) between cargo molecules and MinD protein fluxes (***j***_*p*_ ≠ 0), cargo molecules are not only redistributed due to entropic unmixing effects, but in addition transported along protein gradients by these nonequilibrium protein fluxes (Fig. 2a). As consequence of this additional bias, cargo redistribution is significantly stronger than by equilibrium thermodynamic forces alone, which quantitatively explains our experimental data (Fig. 2b,c). We expect that the coupling constant *ζ*_*pc*_ between MinD and a specific cargo has a small contribution from the origami scaffold as well as from each of its *n* streptavidin: *ζ*_*pc*_ = *ζ*_*po*_ + *n ζ*_*ps*_. This implies that cargo transport should increase with the number of streptavidin integrated into the cargo, as observed in our experiments (Fig. 1d).

## Reduced model predicts protein density-dependent cargo diffusion coefficient

To further elucidate the mechanism underlying MinDE-induced transport, we simplified our theoretical model. Specifically, we neglected membrane saturation effects (see SI for details), so that the chemical potential of a particle with size *a*_*i*_ reduces to *μ*_*i*_ ≈ *k*_*B*_*T* ln(*a*_*i*_*c*_*i*_). Then, the effective force-balance equation, Eq. (1), takes the form of a generalized Fick’s law for the protein fluxes in the non-equilibrium steady state with a density-dependent diffusion coefficient *D*_*p*_(*c*_*c*_, *c*_*s*_):

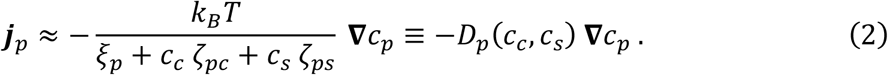

The effective diffusion coefficient *D*_*p*_(*c*_*c*_, *c*_*s*_) decreases upon adding cargo and streptavidin as additional sources of friction. In the presence of cargo with stronger coupling *ζ*_*pc*_, maintaining the diffusive fluxes that balance reactive protein turnover requires sharper protein gradients, which explains our observation of progressively narrower MinDE patterns (Fig. 1f).

Because the number of free streptavidin is typically small (see SI), we assumed that free streptavidin molecules do not significantly contribute to the dynamics, *c*_*s*_ *ζ*_*ps*_ ≪ *c*_*c*_ *ζ*_*pc*_. Using this simplification, we obtain a closed analytical solution for the cargo distribution:

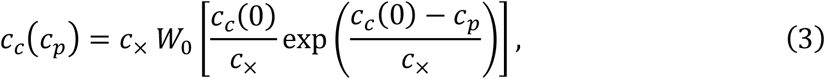

where we defined the *interaction density c*_×_ = *ξ*_*p*_/*ζ*_*pc*_. After fitting Eq. 3 to our experimental data (Fig. 3a), we found—as expected—that the coupling constant *ζ*_*pc*_ increases linearly with the streptavidin count of the cargo (Fig. 3b).

**Fig. 3:**
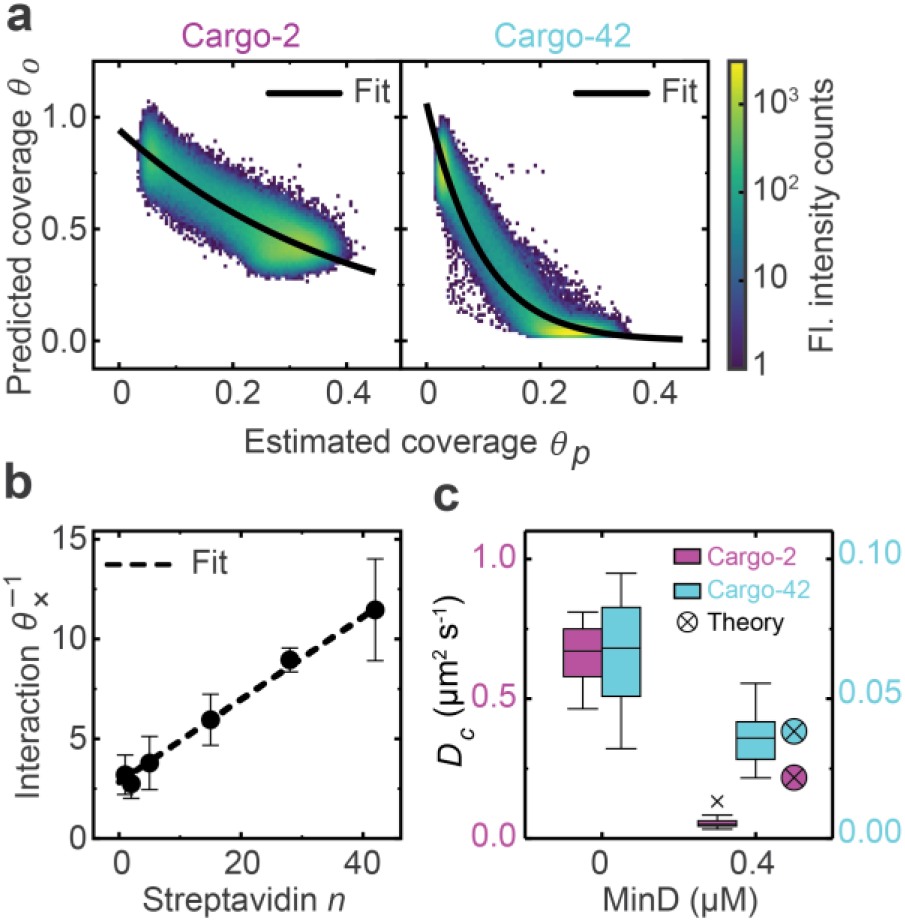
Reduced model predicts slower diffusion at larger ambient MinD density. **a**, Cross-correlation function between MinD coverage (*θ*_*m*_) and cargo coverage (*θ*_*c*_) for cargo-2 and cargo-42. The color-coded 2D-Histogram represents our experimental data, while the solid line is a fit curve of our reduced model. **b**, Interaction parameter (in terms of MinD coverage; surface coverages *θ* = *a c* and surface densities *c* are connected by the particle size *a*) as obtained from our fitting procedure. The interaction linearly increases with the number of streptavidin starting from a basal value (base interaction of the origami scaffold). We find that significant cargo gradients readily build up with a small interaction parameter. However, in contrast to our initial estimate, we have now neglected entropic repulsion between cargo molecules for simplicity. Therefore, we likely underestimate the interaction parameter here (compare to Fig. 2c). **c**, Cargo molecule diffusion coefficient in absence or presence of MinD, as obtained from single-particle tracking and predicted with the fitted interaction parameters obtained in b. Cargo-42 typically diffuses slower than cargo-2 (indicated by a smaller diffusion coefficient *D*_*c*_ at 0 μM MinD). The increase of MinD density has a much stronger effect on cargo-2 than on cargo-42 both in theory and experiment. Box plot lines are median, box limits are quartiles 1 and 3, whiskers are 1.5× interquartile range (IQR) and points are outliers. Data obtained from number of independent experiments E(Cargo-2)=5, E(Cargo-42)=3, E(Cargo-2, MinD)=3, E(Cargo-42, MinD)=2; number of analyzed single particle tracks N(Cargo-2)= 15755, N(Cargo-42)=19481, N(Cargo-2, MinD)=7924, N(Cargo-42, MinD)=4542; average track-length TL(Cargo-2)=339, TL(Cargo-42)=546, TL(Cargo-2, MinD)=772, TL(Cargo-42, MinD)=647; fraction of mobile DNA origami MF(Cargo-2)=0.81, MF(Cargo-42)=0.67, MF(Cargo-2, MinD)=0.70, MF(Cargo-42, MinD)=0.63.

Next, we set out to further test our model experimentally. We performed single particle tracking of two different cargo species, cargo-2 and cargo-42, at low and high MinD densities. In the latter case, we mimicked MinD-rich conditions in MinDE patterns in a simplified fashion by adding 1 μM MinD and ATP, but no MinE. We found that the diffusion coefficient of cargo-2 decreased from 0.65±0.12 μm^2^ s^−1^ in the absence of MinD proteins to 0.06±0.02 μm^2^ s^−1^ at high protein density (Fig. 3c). In contrast, the diffusion coefficient of cargo-42 which was already low in the absence of MinD, 0.06±0.02 μm^2^ s^−1^, hardly changed in the presence of proteins, 0.036±0.011 μm^2^ s^−1^ (Fig. 3c). Subsequently we used our fitted interaction parameters to predict the diffusion coefficient of cargo at high protein densities, based on the experimental values in the absence of proteins (see SI). We found that our predictions were in good agreement with our experimental findings, validating our model. The at first counterintuitive observation that MinD affects cargo-42 diffusion less than that of cargo-2, although the frictional coupling is stronger, can be explained as follows: in this case the friction of the many streptavidin with the membrane becomes dominant compared to the additional friction with MinD. In conclusion, the dependence of the cargo diffusion coefficient on the ambient protein density is a direct experimental proof that friction occurs between MinD and cargo.

## MinDE spatially sort different cargo species

Can we use our obtained knowledge to selectively position cargo molecules, i.e. sort them according to their properties, along protein gradients? To test this idea, we labeled two distinct cargo species, cargo-2 and cargo-42, with different dyes and placed them in the same assay (Fig. 4a). As predicted by our model (Fig. 4d), we found that cargo-42 gathered in MinD-free regions, and were framed by cargo-2 (Fig. 4b, c, Fig. S6, Supplementary Movie 3). Thus, cargo-42 exhibited a similar behavior as when present in the assay alone. In contrast, the localization of cargo-2 in reference to MinD changes when cargo-42 is also present in the assay (compare Fig. 4c and Fig. 1c). The observed spatial separation of cargo species in the established MinDE patterns was not an artefact due to fluorescent channel crosstalk or dye quenching, as it could also be observed when dyes were swapped on the cargo species (Fig. S7). It was also not an artefact caused by the specific dyes used for labelling, since when otherwise identical cargo (either both cargo-2 or cargo-42) were labelled differentially, no spatial sorting could be observed, and the resulting cargo distributions were superimposable (Fig. S8). Hence, the clear MinDE-induced spatial sorting of cargo species according to their effective size further refutes thermodynamic models (Fig. 4d), corroborating that indeed MinDE transport molecules via friction.

**Fig. 4:**
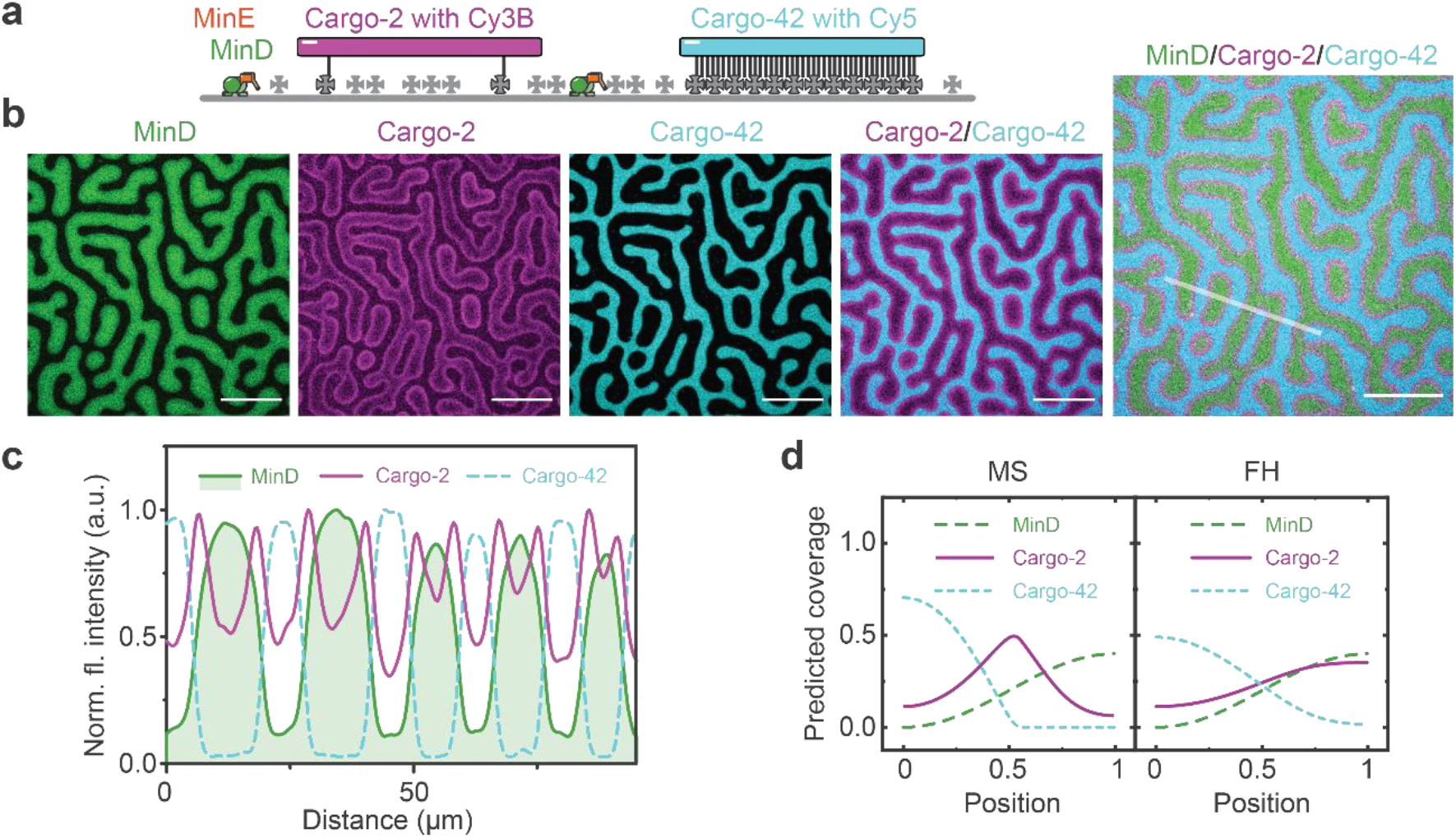
MinDE spatially sorts cargo according to effective size. **a**, Schematic of the experimental setup. MinDE self-organization was performed in presence of two different cargo species with distinct fluorescent labels, cargo-2 with Cy3B and cargo-42 with Cy5 (1 μM MinD (30% EGFP-MinD), 1.5 μM MinE-His, 50 pM origami-Cy3b with 2, and 50 pM origami-Cy5 with 42 biotinylated oligonucleotides, non-labeled streptavidin). **b**, Representative images of individual and overlayed channels, and **c** line plot of indicated selection of MinDE-induced sorting of cargo species. Scale bars: 50 μm. Experiment was performed three times under identical conditions. **d**, Spatial distribution of two cargo species in response to the (imposed) MinD profile, corresponding to the cross-correlation functions in b. The Maxwell-Stefan type model allows for stronger reorganization of cargo molecules than the Flory-Huggins type model. In particular, the Maxwell-Stefan type model predicts that cargo-2 accumulates between cargo-42 and MinD. Model parameters: average coverage of MinD proteins 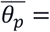 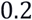, streptavidin 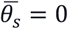, cargo-2 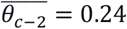 and cargo-42 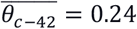; interaction parameter (in terms of MinD coverage) of cargo-2 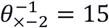 and cargo-42 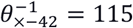. Surface coverages *θ* = *a c* and surface densities *c* are connected by the particle size *a*.

## Conclusion

In conclusion, combining *in vitro* reconstitution experiments with theoretical modeling and analysis, we showed that the prokaryotic MinDE proteins can non-specifically transport and even sort large, membrane-bound cargo molecules by Maxwell-Stefan diffusiophoresis, a process driven by gradient formation of ATP-consuming, active proteins. This constitutes a so far unknown mechanism of coupling energy dissipation to active spatiotemporal positioning in biological systems. Remarkably, we found that the diffusive fluxes of MinD and cargo couple via friction and thus, mechanical rather than thermodynamic forces (Fig. 5). Specifically, active MinDE self-organization generates a net diffusive flux of MinD (towards low MinD densities), which establishes an effective frictional force, driving the accumulation of cargo in areas of low MinD density. Hence, cargoes with fewer streptavidin, i.e. with smaller effective sizes, experience less friction when diffusing through the carpet of MinD molecules on the membrane than those with many streptavidin, i.e. effectively larger ones. Similar transport effects, where concentration gradients of solutes drive diffusiophoretic particle motion, have previously been reported in a non-biological context. For example, they have been shown to emerge from an interaction potential between a large colloidal particle and small solute molecules, which led to fluid slip at the colloid surface^32,37,38^ or were driven by electrostatic interactions^34–36^. In addition, particle fluxes can also couple directly via friction, which has been described as Maxwell-Stefan diffusion^28,29^ and experimentally demonstrated for three-component gases^30,31^. In a biologically relevant setting, so far only diffusiophoresis of large particles driven by gradients of small biomolecules such as ATP has been theoretically suggested^38^. In contrast, Maxwell-Stefan diffusiophoresis has, to the best of our knowledge, not yet been described to drive molecular transport in biology. In comparison with translational motor proteins known from higher organisms, this mechanism could be interpreted as an alternative, more rudimentary mode of mechanochemical coupling. This mechanism is presumably not a special feature of the *E. coli* MinDE system or reaction-diffusion systems in general, but can potentially be exerted by any active system that generates and maintains concentration gradients. For example, such a mechanism could be at play for the plethora of intracellular (actin) waves that have been discovered in eukaryotes and whose purpose and mode of action has remained elusive^45^. The mechanism might not even be limited to the membrane as a reaction surface, but potentially extends to other cellular surfaces and even cytosolic gradients. For instance, the strong concentration gradients that are built up during liquid-liquid phase separation processes could potentially impact other molecules in a similar fashion^46^. From the results obtained here, we can also predict that distinct pattern-forming systems that share the same reaction space should typically align to minimize friction, even if their constituents are chemically independent. This could potentially link and synchronize multiple pathways in a simple fashion to increase their efficacy or provide a rescue mechanism against mutations affecting the chemical coupling via specific interactions (for example between MinC and FtsZ). That this non-specific means of transport was discovered and described in detail in an *in vitro* reconstitution assay is not a coincidence, but highlights that the complexity of cells with more sophisticated and stronger specific interactions presumably masks such occurrence. Lastly, simple as it is, this mechanism might be prevalent in prokaryotes and constitute an archetypal mechanism present in early forms of life.

**Fig. 5:**
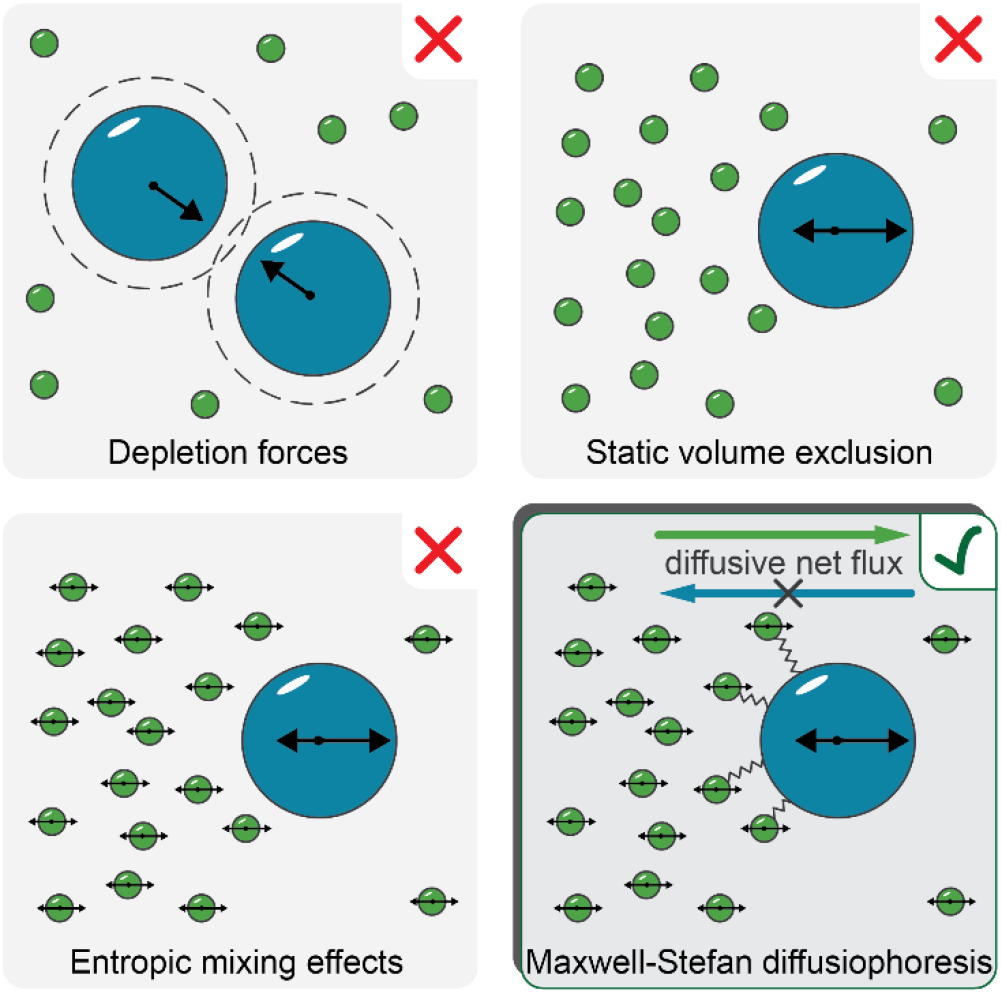
Maxwell-Stefan diffusiophoresis explains cargo transport by protein self-organization. Thermodynamic forces, such as depletion forces, static volume exclusion or entropic mixing effects cannot explain MinDE-dependent cargo transport. However, it can be explained by Maxwell-Stefan diffusiophoresis: active protein self-organization generates gradients and hence net diffusive fluxes. As the proteins interact non-specifically with the cargo, the diffusive fluxes establish an effective frictional force, driving the accumulation of cargo in areas of low protein density.

## Methods

### Plasmids and proteins

The plasmids pET28a-His-MinD_MinE^13^, pET28a-His-EGFP-MinD^47^ and pET28a-MinE-His^15^ were used for purification of His-MinD, His-EGFP-MinD and MinE-His, respectively, as described in detail previously^48^. In brief, proteins were expressed in *E. coli* BL21 (DE3) and then purified via Ni-NTA affinity and size-exclusion chromatography in storage buffer (50 mM HEPES/KOH pH 7.2, 150 mM KCl, 10% glycerol, 0.1 mM EDTA, 0.4 mM TCEP). Proteins were snap-frozen in liquid nitrogen and stored in small aliquots until further use at −80 °C.

### DNA origami nanostructures

The elongated DNA origami nanostructure used here has been previously designed and described^39^. The 20-helix bundle with hexagonal lattice is based on the M13mp18 7429-nucleotide long scaffold plasmid (p7429) (Bayou Biolabs, Metairie, LA, USA), and was modified using CaDNAno^49^. Staple oligonucleotides, 5’-Cy3B/Cy5-functionalized oligonucleotides oligonucleotides (Both: High purity salt free, Eurofins MWG Operon, Ebersberg, Germany) and 5’-biotin-TEG functionalized (Sigma-Aldrich, St. Louis, USA) were purchased or diluted in Milli-Q ultrapure water at a concentration of 100 μM. Origami structures with 1-15 anchors were based on the previous design^39^, which was further modified for functionalization with 42 anchors (Fig. S9). The assembly of the origami structure was performed in a one-pot reaction mix as described previously^39^. In brief, the components were mixed at a final concentration of 20 nM p7429 scaffold plasmid and 200 nM staple oligonucleotides in folding buffer (5 mM Tris-HCl, 1 mM EDTA, 20 mM MgCl_2_, pH 8.0) and annealed in a thermocycler (Mastercycler, Eppendorf, Hamburg, Germany) over a 41 h cooling scheme from 65 to 40 °C. Folded nanostructures were purified to remove excess staple strands by centrifugation (14000 g, three cycles for 3 min, one cycle for 5 min) in Amicon Ultra 100 kDa MWCO filters (Merck Millipore, Darmstadt, Germany) using reaction buffer (25 mM Tris-HCl pH 7.5, 150 mM KCl, 5 mM MgCl_2_). The concentration of folded Cy5-labeled origami structures was estimated by fluorescence intensity measurements using a one-drop measurement unit of a Jasco FP-8500 spectrofluorometer (Tokyo, Japan) and subsequent comparison with an intensity calibration curve obtained for free Cy5 dye corrected for the multiple labelling of the origami. Cy3B-labeled DNA origami concentration was measured by absorption at 260 nm on a Nanodrop Spectrophotometer (ThermoFisher Scientific, Waltham, USA) and related to Cy5-labeled structures of known concentrations. Cy3B/Cy5-labeled DNA origami structures contained 7 Cy3B/Cy5-labelled oligonucleotides attached to extended staples on the upper facet. At the lower facet the origami contained multiple 18 nt extensions that were hybridized with complementary 5’Biotin-TEG-functionalized oligonucleotides at defined positions.

### Preparation of supported lipid bilayers

Supported lipid bilayers (SLBs) were prepared as described in detail before^13,48^. In brief, cover slides were rinsed with ddH_2_O and ethanol, and a plastic chamber was glued on top. Slides were further cleaned by plasma cleaning with oxygen as process gas (model Zepto, Diener electronic, Ebhausen, Germany). Chloroform-dissolved lipids (Avanti Polar Lipids, Alabaster, AL, USA) were dried by a nitrogen stream and subsequently in a desiccator before slow rehydration at a concentration of 4 mg/ml in reaction buffer (25 mM Tris-HCl pH 7.5, 150 mM KCl, 5 mM MgCl_2_). Small unilamellar vesicles were generated by sonication in a bath sonicator and subsequently added to the cleaned reaction chambers. All mentioned concentrations refer to the final volume of the reaction chamber of 200 μl. To prepare chambers for self-organization experiments the SLB was generated with a lipid composition of 69/30/1 mol% DOPC/DOPG/Biotinyl-CAP-PE or with 70/30/0.01 mol% DOPC/DOPG/Biotinyl-CAP-PE for single particle tracking experiments and subsequently incubated with non-labelled or Alexa568-labelled streptavidin (ThermoFisher Scientific, Waltham, USA) at a final concentration of 1 μg/ml. After 5-10 min incubation unbound streptavidin was removed by washing 5 times with a total volume of 1 ml reaction buffer. The buffer was adjusted to 100 μl volume and the origami was incubated at a final concentration of 0.1 nM for 10 min, before the buffer was adjusted to the final volume of 200 μl. For experiments involving more than one type of DNA origami, DNA origami species were premixed in DNA LoBind tubes (Eppendorf, Hamburg, Germany) before addition to the sample chamber at a final concentration of 50 pM for each DNA origami, keeping the overall DNA origami concentration at 0.1 nM. For single particle tracking experiments DNA origami was diluted in DNA LoBind tubes (Eppendorf) and added to a chamber at a final concentration of 0.1-1 pM. Note that at these experimental conditions, DNA origami does not bind non-specifically to the lipid membrane in the absence of Biotin-TEG-anchors or streptavidin, due to the high net negative charge of both SLB and DNA origami^50,51^.

### Single particle tracking

Single particle tracking of DNA origami was conducted at a concentration of DNA origami and anchors that can be described as diluted, so that interaction between individual DNA origami was minimized^52^ (SLB: 70/30/0.01 mol% DOPC/DOPG/Biotinyl-CAP-PE, 0.1-1 pM origami). Due to the superior brightness and photostability, single particle tracking was exclusively performed using Cy3B-labeled DNA origami. To further reduce photobleaching and blinking as well photopolymerization of MinD single particle tracking was performed in the presence of an oxygen scavenger system (3.7 U/ml pyranose oxidase, 90 U/ml catalase, 0.8% glucose)^53^ and trolox.

### Microscopy

All images, except for single-particle tracking, were taken on a Zeiss LSM780 confocal laser scanning microscope using a Zeiss C-Apochromat 40×/1.20 water-immersion objective (Carl Zeiss AG, Oberkochen, Germany). Longer time-series were acquired using the built-in autofocus system. All two or three color images were acquired with alternating illumination for the 488/633 nm and 561 nm laser lines to avoid cross-talk. EGFP-MinD was excited using the 488 nm Argon laser, Cy3B-labeled origami or Alexa568-streptavidin using the 561 nm DPSS laser and Cy5-labeled origami using the 633 nm He–Ne laser. Images were typically recorded with a pinhole size of 2.6-4 Airy unit for the EGFP and origami channels, and 1 Airy unit for the streptavidin channel, 512×512 pixel resolution, and a pixel dwell time of 1.27 μs. Time-series were typically acquired with about 14 s intervals. For single particle tracking of DNA origami images were acquired on a custom-built total internal reflection fluorescence microscope (TIRFM)^54^ using a NIKON SR Apo TIRF 100×/1.49 oil-immersion objective, constructed around a Nikon Ti-S microscope body (both Nikon GmbH, Düsseldorf, Germany). Two laser lines (490 nm (Cobolt Calypso, 50mW nominal) and 561 nm (Cobolt Jive, 50mWnominal), Cobolt AB, Solna, Sweden)) were controlled in power and timing (AOTF, Gooch&Housego TF-525-250, Illminster, UK) and spatially filtered (kineFLEX-P-3-S-405.640-0.7-FCS-P0, Qioptiq, Hamble, UK). The beam was further collimated, expanded (10x) and focused on the objective’s back aperture by standard achromatic doublet lenses. The TIRF angle was controlled by precise parallel offset of the excitation beam (Q545, PI, Karlsruhe, Germany). The emission light was notch filtered to remove residual excitation light, spectrally separated by a diochroic beamsplitter (T555lpxr-UF1, Chroma Technology Cooperation, Bellow Falls, VT), bandpass filtered, 525/50 and 593/46 (both Chroma), respectively, and re-positioned on two halves of the EMCCD camera (Andor iXon Ultra 897, Andor Technologies, Belfast, UK). Images were recorded with Andor Solis (Ver. 4.28, Andor Technologies).

### Image Analysis

All images were processed using Fiji (version v1.52p), Matlab (R2016a, The Math-Works, Natick, USA) or Python (Python Software Foundation). Brightness or contrast adjustments of all displayed images were applied homogenously.

For line plots images were smoothed with a Gaussian filter with pixel width 2 in Fiji. Images were smoothed with a Gaussian kernel of 1 pixel width. The theoretical models were formulated as boundary-value problems and solved in a 1D geometry using a finite-difference scheme using SciPy^55^. Curve fitting was performed with lmfit^56^.

### Single Particle Tracking Analysis

Analysis of single particle tracking was conducted as described previously using previously established code^57^. In brief, a custom-written MATLAB code was used to detect DNA origami fluorescence in each frame and extract their position. Origami trajectories on the membrane were analyzed using jump-distance (JD) analysis^58,59^. The distances between particle locations between subsequent frames were analyzed and diffusion coefficients of particle ensembles were obtained by fitting the cumulative histograms. As usually some of the origami in the field of view were immobile and did not diffuse, cumulative histograms of obtained jump distances were fitted with 2 components, where for the second component the upper boundary was set to 0.1 μm^2^ s^−1^, and usually resulted in diffusion coefficients of less than 0.01 μm^2^ s^−1^.

### Analysis of MinDE-Dependent Transport

Analysis of fluorescence intensities and contrast was essentially performed as described earlier^7^. In brief, tile scans were imported into Fiji, where the EGFP-MinD channel was used for segmentation to generate a binary mask of the patterns. The original non-modified images from the two/three spectral channels were analyzed based on the binary mask using a custom-written Matlab code. The average fluorescence intensity in the Alexa568-streptavidin or origami-Cy5 and EGFP-MinD spectral channel was obtained by pooling the means of individual images from one independent experiment. All means from one independent experiment and condition were pooled together. All fluorescence intensity values from one experimental set were normalized to the fluorescence intensity values obtained for the respective origami with 1 anchor. The contrast of the resulting patterns was calculated for every individual image as the difference between the average intensity in the MinDE minima and MinDE maxima divided by the average intensity in the MinDE maxima. The contrast of the MinDE patterns was calculated for every individual image as the difference between the average intensity in the MinDE maxima and MinDE minima divided by the average intensity in the MinDE minima.

## Supporting information

Description Supplementary Movies

## Acknowledgements

We thank MPIB Core Facility for assistance in protein purification. We further thank Hiromune Eto and Henri Franquelim for helpful discussions. B.R., E.F. and P.S. acknowledge funding by the German Research Foundation (DFG) – Project 269423233 – TRR 174. B.R., A.K. and A.G. were supported by a DFG fellowship through the Graduate School of Quantitative Biosciences Munich (QBM). P.S. acknowledges the support of the research network MaxSynBio via a joint funding initiative of the German Federal Ministry of Education and Research (BMBF) and the Max Planck Society.

## Author contributions

BR, AG, EF and PS conceived the study. AG and EF designed the theoretical analysis. AG conducted the theoretical analysis. BR, AK and PB designed experiments. BR performed all experiments. AK designed DNA origami. AK and BR prepared origami. KG developed single-particle tracking code. BR, AG, PB and KG analysed data. BR, AG, EF and PS wrote the paper. All authors discussed and interpreted results and revised the manuscript.

