## Supplementary material for "ATP driven diffusiophoresis: active cargo transport without motor proteins": Description Supplementary Movies

**Supplementary Movie 1**: Representative time-series of MinDE self-organization inducing patterns of DNA origami and streptavidin when no origami, cargo-2, cargo-15 or cargo-42 is present (1 µM MinD (30% EGFP-MinD), 1.5 µM MinE-His, in absence or presence of 0.1 nM origami-Cy5 with 2, 15 or 42 biotinylated oligonucleotides, Alexa568-streptavidin). Scale bars: 50 µm

**Supplementary Movie 2:** Representative time-series and kymograph showing changes to MinD and cargo molecule patterns in presence of cargo-1, cargo-2, cargo-15 or cargo-42 (1 µM MinD (30% EGFP-MinD), 1.5 µM MinE-His, origami-Cy5 with 1, 2, 15 or 42 biotinylated oligonucleotides, Alexa568-streptavidin) upon addition of more MinE (addition of 1.5 µM MinE-His). MinE addition directly before t = 0 s. Scale bars: 50 µm

**Supplementary Movie 3**: Representative time-series of MinD, cargo-2 and cargo-42 pattern formation. ATP is added to start self-organization directly before t=0 s (1 µM MinD (30% EGFP-MinD), 1.5 µM MinE-His, 50 pM origami-Cy3B with 2 biotinylated oligonucleotides, 50 pM origami-Cy5 with 42 biotinylated oligonucleotides, non-labelled streptavidin). Scale bars: 50 µm
