## Supplementary Information for "ATP driven diffusiophoresis: active cargo transport without motor proteins"

### Supplementary Theoretical Description

#### I. Chemical potential gradients induce flows, which couple by an out-of-equilibrium Maxwell-Stefan mechanism

Elaborating the brief description in the main text, we here explain our theory in full detail. First, we comment on the abundance of membrane-bound molecules, which is a necessary information to formulate our theory. Then, we describe the population of membrane-bound molecules as a lattice gas. Specifically, we use the Flory-Huggins theory of mixing to calculate the local chemical potential of each species. Finally, we consider chemical-potential-induced particle flows, and their coupling via an effective inter-particle friction (Maxwell-Stefan approach to diffusion).

##### 1. Cargo density is limited by abundance of streptavidin and origami

Each cargo molecule is a composite object consisting of a DNA origami scaffold and multiple streptavidin building blocks. Therefore, the total number of membrane-bound cargo ($N_c$) should be limited by the abundance of DNA origami ($N_{o+}$) and membrane-bound streptavidin ($N_s$). Because of their strong binding, we expect that all available streptavidin molecules will bind to biotinylated lipids that are incorporated into the membrane. Thus, the average density of streptavidin molecules on a membrane of size $A$ is given by $\overline{c_s} = N_s/A$. Each DNA origami scaffold has $n$ biotinylated oligonucleotide handles that can attach to membrane-bound streptavidin. If all DNA origami were to bind to the membrane, then this would correspond to an average density of $\overline{c_{o+}} = N_{o+}/A$. For small numbers of biotinylated oligonucleotide handles per origami, this leaves some freely diffusing (not bound to DNA origami scaffolds) streptavidin molecules on the membrane.

In contrast, for large numbers of biotinylated oligonucleotide handles per origami, all streptavidin molecules will bind to DNA origami scaffolds. Then, the DNA origami scaffolds will compete for the available membrane-bound streptavidin molecules, effectively resulting in an average membrane-bound cargo density of $\overline{c_c} = N_c/A$, while leaving $N_{o+} - N_c$ DNA origami scaffolds unbound. In our experiments, membrane-bound DNA origami scaffolds cover up to 60% of the membrane, resulting in a strong steric repulsion due to volume exclusion effects. Thus, we expect that each individual DNA origami scaffold with $n$ biotinylated oligonucleotide handles will maximize its adhesion to the membrane by binding exactly $n$

streptavidin building blocks (Fig. 2a). Therefore, the maximal abundance (average concentration) of membrane-bound cargo is given by $\overline{c_c} = \min(\overline{c_{o+}}, \overline{c_s}/n)$.

With the given amount of streptavidin on the membrane and origami in the assay (Table S1), we would expect that all streptavidin molecules are bound to DNA origami scaffolds for $n \geq \overline{c_s}/\overline{c_{o+}} = 18.75$. Consequently, for origami with larger number of streptavidin, (n=28 and 42), not all origami can bind to membrane-bound streptavidin, decreasing their overall membrane density. In good agreement with these arguments, the average fluorescence intensity of DNA origami scaffolds in our experiments indicates that the average density of membrane-bound streptavidin molecules is the limiting factor for binding of DNA origami to the membrane for $n \geq 15$ (Fig. S5b).

#### 2. Flory-Huggins theory of mixing: an equilibrium picture

Because our setup resembles polymers in solution, we formulated a Flory-Huggins theory by tessellating the membrane into (infinitesimally) small well-mixed compartments. Specifically, we asked: given a heterogeneous distribution of active particles (i.e., we allow MinD protein density gradients across adjacent compartments), what is the equilibrium distribution of passive particles (cargo and free streptavidin molecules)? To answer this question, we derived the corresponding chemical potentials $\mu_i$ for each species as discussed next.

Before formulating a model, we took a closer look at the microscopic geometry of the cargo molecules in our experiments (see illustration "Conceptualized model geometry"). Each cargo molecule is a composite object which consists of a DNA origami scaffold and multiple streptavidin building blocks. The rod-shaped DNA origami resides between 5 and 11 nm above the membrane (see Supplementary Note 1). The streptavidin building blocks also serve as membrane tethers with a height of roughly 5 nm. Furthermore, MinD proteins bind to the membrane in a monomolecular layer of about 5 nm height[1]. Therefore, we expect that cargo transport is dominated by interactions between MinD and streptavidin. The geometry of our cargo molecules signifies two distinct interaction layers, which we indicate with the following labels (see illustration "Conceptualized model geometry"): ($\sigma$) the *proximal plane* refers to the thin layer near the membrane, which has a height of 11 nm, and ($\tau$) the *distal plane* refers to the thin layer above the proximal plane, which has a height of 8 nm. We regarded these two layers, the proximal plane and the distal plane, as two distinct lattice gases that are linked by the cargo molecules (which are present in *both* layers with a common local density $c_c$ but

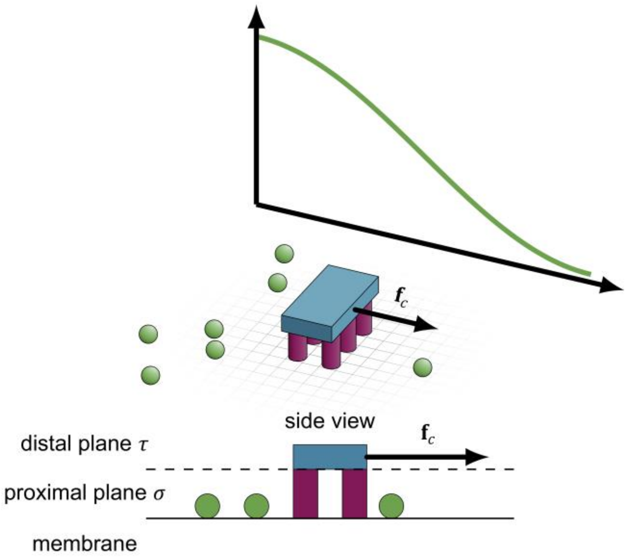

**Conceptualized model geometry (cf. Fig. 2a in the main text).** Cargo molecules are composite objects which consist of a DNA origami scaffold (blue) and multiple streptavidin building blocks (magenta). The rod-shaped DNA origami resides between 5 and 11 nm above the membrane (see Supplementary Note 1). The streptavidin building blocks also serve as membrane tethers. Furthermore, MinD proteins (green spheres) bind to the membrane in a monomolecular layer of about 5 nm height[1]. Therefore, we expect that cargo transport is dominated by interactions between MinD and streptavidin. This setup signifies two distinct interaction layers of the cargo with the proteins, which we indicate with the following labels: ($\sigma$) In the *proximal plane*, each DNA origami crosslinks multiple streptavidin blocks into a polymer-like object *c*, which mingles in a sea of MinD proteins *p*, free streptavidin molecules *s*, and solvent. ($\tau$) In the *distal plane*, we assume that only solvent surrounds the bulky DNA origami body *o*. We describe both interaction layers as lattice gases (indicated by the black grid), but with different lattice constants in the proximal and in the distal plane. In the Maxwell-Stefan picture, MinDE protein density gradients lead to diffusive flows (green arrows), which exert an effective force $\mathbf{f}_c$ on the cargo molecule (black arrow).

distinct effective sizes $a_c$ and $a_o$). We choose the lattice constants of the proximal plane and the distal plane to match the typical length of the associated particles, respectively.

On the membrane (*proximal plane* $\sigma$), each DNA origami scaffold crosslinks multiple streptavidin blocks into a polymer-like object $c$, which mingles in a carpet of MinD proteins $p$, free streptavidin molecules $s$, and solvent. Because streptavidin and MinD proteins have a diameter of roughly 5 nm, we choose the lattice constant of the proximal plane accordingly: $\ell_\sigma = 5\,\text{nm}$. Then, each individual protein, free streptavidin molecule, and patch of solute occupies one lattice site of size $a_\sigma = \ell_\sigma^2$. Cargo molecules with $n$ streptavidin building blocks occupy $n$ lattice sites and therefore an effective size $a_c = n\, a_\sigma$ on the membrane. We assume that these four species of molecules form a lattice gas with a free energy density $f^\sigma$, which is described by the Flory-Huggins theory of mixing[2–4]:

$$\frac{f^\sigma}{k_B T} = c_i \ln \theta_i + \; c_i \theta_j \, \chi_{ij}. \tag{S1}$$

Here, $c_i$ and $\theta_i$ refer to the surface density and surface coverage of each species (cargo $c$, free streptavidin $s$, proteins $p$ and solvent $\varrho$), respectively, while the parameters $\chi_{ij}$ are a measure for the interaction energy between different species. We assume that there is no direct attraction or repulsion between the different species (e.g. due to electrostatic interactions); thus, all six possible interaction parameters between the different species vanish: $\chi_{ij} = 0$. As we have tessellated the membrane into (infinitesimally) small well-mixed compartments, we assume that each of these individual compartments in a local equilibrium. Then, by taking the derivative of the free energy density (Eq. S1) with respect to the surface density of cargo molecules, we find the contribution of the *proximal plane* $\sigma$ (near the membrane) to the total chemical potential of a cargo molecule:

$$\frac{\mu_c^\sigma(\boldsymbol{x})}{k_B T} = \frac{\partial}{\partial c_c} \frac{f^\sigma}{k_B T} = \ln[\theta_c(\boldsymbol{x})] + n \ln\left[1 - \theta_c(\boldsymbol{x}) - \theta_s(\boldsymbol{x}) - \theta_p(\boldsymbol{x})\right] + (1 - n). \tag{S2a}$$

Here, $\theta_c(\boldsymbol{x}) = n \, a_\sigma \, c_c(\boldsymbol{x})$ indicates the local surface fraction occupied by cargo molecules, $\theta_s(\boldsymbol{x}) = a_\sigma \, c_s(\boldsymbol{x})$ indicates the local surface fraction occupied by free streptavidin molecules, and $\theta_p(\boldsymbol{x}) = a_\sigma \, c_p(\boldsymbol{x})$ indicates the surface fraction occupied by proteins. Analogously, the chemical potential of a free streptavidin molecule is given by

$$\frac{\mu_s^\sigma(\boldsymbol{x})}{k_B T} = \frac{\partial}{\partial c_s} \frac{f^\sigma}{k_B T} = \ln[\theta_s(\boldsymbol{x})] + \ln\left[1 - \theta_c(\boldsymbol{x}) - \theta_s(\boldsymbol{x}) - \theta_p(\boldsymbol{x})\right], \tag{S2b}$$

and the chemical potential of a membrane-bound MinD protein is given by

$$\frac{\mu_p^\sigma(\boldsymbol{x})}{k_B T} = \frac{\partial}{\partial c_p} \frac{f^\sigma}{k_B T} = \ln\left[\theta_p(\boldsymbol{x})\right] + \ln\left[1 - \theta_c(\boldsymbol{x}) - \theta_s(\boldsymbol{x}) - \theta_p(\boldsymbol{x})\right]. \tag{S2c}$$

At an altitude of 11 nm above the membrane, the *distal plane* $\tau$ contains only DNA origami scaffolds and solvent. Because each DNA origami scaffold is 16 nm wide and 110 nm long, we choose the lattice constant of the proximal plane accordingly: $\ell_\tau = 16$ nm. Then, each DNA origami scaffold occupies roughly seven lattice sites with a total size of $a_o = 7 \, a_\tau$ (yielding a width of 16 nm and a length of 112 nm), while each patch of solute has an effective size of $a_\tau = \ell_\tau^2$. Analogously to Eq. S1, we assume that these two species of molecules form a lattice gas with a free energy density $f^\tau$, which is described by the Flory-Huggins theory of mixing[2–4]:

$$\frac{f^\tau}{k_B T} = c_i \ln \theta_i + \; c_i \theta_j \, \chi_{ij}. \tag{S3a}$$

Here, $c_i$ and $\theta_i$ refer to the surface density and surface coverage of each species (cargo $c$ and solvent $\varrho$), respectively, while $\chi_{c\varrho}$ is a measure for the interaction energy between the two species. As before, we assume that there is no direct attraction or repulsion between the different species (e.g. due to electrostatic interactions); thus, the possible interaction parameter between the two species vanishes: $\chi_{c\varrho} = 0$. As we have tessellated the membrane into (infinitesimally) small well-mixed compartments, we assume that each of these individual compartments is in a local equilibrium. Then, by taking the derivative of the free energy density (Eq. S3a) with respect to the surface density of cargo molecules, we find the contribution of the *distal plane* $\tau$ (11 nm above the membrane) to the total chemical potential of a cargo molecule:

$$\frac{\mu_c^\tau(\boldsymbol{x})}{k_B T} = \frac{\partial}{\partial c_c} \frac{f^\tau}{k_B T} = \ln[\theta_o(\boldsymbol{x})] + 7 \ln[1 - \theta_o(\boldsymbol{x})] - 6. \tag{S3b}$$

Here, $\theta_o(\boldsymbol{x}) = 7 \, a_\tau \, c_c(\boldsymbol{x})$ refers to the surface fraction occupied by the bulky DNA origami scaffolds.

The total chemical potential of a cargo molecule has a contribution from the proximal plane $\sigma$ (Eq. S2a) and a contribution from the distal plane $\tau$ (Eq. S3b), which we weight according to the thickness of the corresponding layer (proximal plane: 11 nm; distal plane: 8 nm). This weighting procedure is equivalent to integrating the free energy density across the thickness of both the proximal plane $\sigma$ (near the membrane) and the distal plane $\tau$ (cf. Eqs. S1 and S3a; the particle densities need to be treated as volumetric densities during the integration and then projected back to surface densities). Then, the total chemical potential of a cargo molecule is given by

$$\mu_c(\boldsymbol{x}) = \frac{11}{19} \mu_c^\sigma(\boldsymbol{x}) + \frac{8}{19} \mu_c^\tau(\boldsymbol{x}). \tag{S3c}$$

In our experimental assay, ATP-consuming reactions continuously drive the protein distribution out of equilibrium and maintain a non-equilibrium steady state with finite protein gradients[5–8]. In stark contrast to MinDE, cargo molecules and free streptavidin molecules are purely passive and almost permanently bound to the membrane, as they have a negligible detachment rate due to their strong membrane anchoring[9]. Therefore, one can assume that the passive particles, cargo $c_c$ and free streptavidin molecules $c_s$, adopt a thermal equilibrium state lacking any gradients in the chemical potentials ($\boldsymbol{\nabla}\mu_c = \boldsymbol{\nabla}\mu_s = 0$; i.e. passive Brownian particles in adjacent compartments have identical chemical potential) in an adiabatic response to the externally imposed distribution of active particles (MinD proteins $c_p$; $\boldsymbol{\nabla}\mu_p \equiv \boldsymbol{\nabla}\mu_p^\sigma \neq 0$). Given these constraints, we then determined the distribution of all passive particles as follows.

The constraint $\boldsymbol{\nabla}\mu_c = 0$ corresponds to the following partial differential equation:

$$\left[\frac{11}{19}\frac{\partial \mu_c^\sigma}{\partial \theta_c}\frac{\partial \theta_c}{\partial \theta_o} + \frac{8}{19}\frac{\partial \mu_c^\tau}{\partial \theta_o}\right]\boldsymbol{\nabla}\theta_o + \frac{11}{19}\frac{\partial \mu_c^\sigma}{\partial \theta_s}\boldsymbol{\nabla}\theta_s + \frac{11}{19}\frac{\partial \mu_c^\sigma}{\partial \theta_p}\boldsymbol{\nabla}\theta_p = 0, \tag{S4a}$$

while the constraint $\boldsymbol{\nabla}\mu_s \equiv \boldsymbol{\nabla}\mu_s^\sigma = 0$ corresponds to:

$$\frac{\partial \mu_s^\sigma}{\partial \theta_c}\frac{\partial \theta_c}{\partial \theta_o}\boldsymbol{\nabla}\theta_o + \frac{\partial \mu_s^\sigma}{\partial \theta_s}\boldsymbol{\nabla}\theta_s + \frac{\partial \mu_s^\sigma}{\partial \theta_p}\boldsymbol{\nabla}\theta_p = 0. \tag{S4b}$$

The ratio $\partial\theta_c/\partial\theta_o = a_c/a_o = (7\ a_\tau)/(n\ a_\sigma)$ is fully determined by the number of streptavidin blocks per cargo. Since the distribution of proteins, i.e. $\boldsymbol{\nabla}\theta_p$, is externally maintained, Eqs. S4a and S4b form a closed set of partial differential equations. Note that neither the precise functional form of the protein coverage distribution $\theta_p$ nor the dimension of the geometry are important, for the following reason: In general, the equilibrium distribution of the cargo coverage will be determined by the distribution of proteins and the abundance of molecules in the assay, and therefore have the form $\theta_o(\theta_p, \overline{\theta_p}, \overline{\theta_o}, \overline{\theta_s}, n)$. Similarly, the equilibrium distribution of the streptavidin coverage has the form $\theta_s(\theta_p, \overline{\theta_p}, \overline{\theta_o}, \overline{\theta_s}, n)$. Thus, by using the chain rule of differentiation, one could in principle fully eliminate all gradients from Eqs. S4a and S4b:

$$\left[\frac{11}{19}\frac{7\ a_\tau}{n\ a_\sigma}\frac{\partial \mu_c^\sigma}{\partial \theta_c} + \frac{8}{19}\frac{\partial \mu_c^\tau}{\partial \theta_o}\right]\frac{\partial \theta_o}{\partial \theta_p} + \frac{11}{19}\frac{\partial \mu_c^\sigma}{\partial \theta_s}\frac{\partial \theta_s}{\partial \theta_p} + \frac{11}{19}\frac{\partial \mu_c^\sigma}{\partial \theta_p} = 0, \tag{S5a}$$

$$\frac{7\ a_\tau}{n\ a_\sigma}\frac{\partial \mu_s^\sigma}{\partial \theta_c}\frac{\partial \theta_o}{\partial \theta_p} + \frac{\partial \mu_s^\sigma}{\partial \theta_s}\frac{\partial \theta_s}{\partial \theta_p} + \frac{\partial \mu_s^\sigma}{\partial \theta_p} = 0, \tag{S5b}$$

and directly solve for the coverage of passive particles as a function of the protein coverage. Note that Eqs. S5a and S5b can also be obtained by directly setting $\mu_c = cst$ and $\mu_s = cst$, and expanding the resulting equations to first order in the protein coverage $\theta_p$. Alternatively, one can also obtain Eqs. S5a and S5b by integrating Eqs. S4a and S4b over an arbitrary infinitesimal line segment $\boldsymbol{n}\,ds$, and perform a change of variables $ds\,\boldsymbol{n}\cdot\boldsymbol{\nabla}\theta = d\theta$.

Since we were also interested in the spatial distribution of passive molecules, however, we translated Eqs. S4a and S4b into a boundary value problem, in a 1D geometry of length $L \equiv 1$.

Specifically, we introduced two fields $\Theta_o = \frac{1}{L}\int_0^x dy\, \theta_o(y)$ and $\Theta_s = \frac{1}{L}\int_0^x dy\, \theta_s(y)$, resulting in the additional two differential equations

$$\theta_o = L\, \nabla\Theta_o, \qquad \text{(S4c)}$$
$$\theta_s = L\, \nabla\Theta_s. \qquad \text{(S4d)}$$

Thus, in summary, we have four partial differential equations S4a-d with the four boundary conditions $\Theta_s(0) = 0$, $\Theta_s(L) = \overline{\theta_s}(n)$ , $\Theta_o(0) = 0$, and $\Theta_o(L) = \overline{\theta_o}(n)$. As we have discussed in section 1 "Cargo density is limited by abundance of streptavidin and origami", the average coverage of free streptavidin molecules and cargo (equivalent to their density or abundance) depend on the number of streptavidin blocks per cargo molecule. Then, we imposed the following coverage distribution of proteins:

$$\theta_p = \overline{\theta_p}\, \frac{2}{L} \cos\left(\frac{\pi\, x}{2\, L}\right)^2, \qquad (S6)$$

and solved the closed set of partial differential equations S4a-d.

We found that entropic mixing effects can in principle lead to transport of passive particles in a gradient of active particles (Fig. 2b,c). However, this disagrees with our experiments where we observed a far stronger redistribution of the passive cargo molecules than entropic mixing would predict (Figure 2b). In other words, the weak entropic sorting of the cargo's small streptavidin blocks in a fixed gradient of MinD proteins is not sufficient to overcome the strong entropic repulsion between the large origami scaffolds: as the area fraction $\theta_o$ that is covered by DNA origami scaffolds approaches saturation, $\theta_o \to 1$, the second term in Eq. S3b diverges logarithmically and prevents further agglomeration.

##### 3. Maxwell-Stefan coupling between diffusive fluxes: an out-of-equilibrium picture

As described in the main text, we next relaxed our previous assumption of fixed external chemical potential gradients and considered their dynamics. According to Onsager's theory of nonequilibrium thermodynamics[10], gradients in a chemical potential $\nabla\mu_i$ imply particle fluxes $\boldsymbol{j}_i$. In the present context, a possible candidate for a non-equilibrium process in a crowded environment is the coupling of particle fluxes through friction caused by non-specific interactions between proteins and cargo molecules on the membrane. Such coupling between diffusive fluxes has been predicted by the Maxwell-Stefan theory of diffusion[11,12] and has been experimentally observed for three-component gas mixtures[13,14]. The theory asserts that each species on the membrane obeys an effective force-balance equation[11,12]:

$$\nabla\mu_i + \xi_i \frac{\boldsymbol{j}_i}{c_i} + \sum_k c_k\,\zeta_{ik}\left(\frac{\boldsymbol{j}_i}{c_i} - \frac{\boldsymbol{j}_k}{c_k}\right) = 0, \qquad \left(\begin{matrix}\text{S7,}\\ \text{1 in main text}\end{matrix}\right)$$

where the index $i \in \{p, c, s\}$ refers to MinD proteins, cargo molecules with bound streptavidin, and free streptavidin molecules, respectively. In the nonequilibrium steady state, thermodynamic driving forces, caused by chemical potential gradients, are balanced by friction forces between the different macromolecules and lipid molecules (with friction coefficient $\xi_i$) as well as among the macromolecules themselves (with coupling constants $\zeta_{ik}$). Note that, according to Onsager's relations[10], the matrix of coupling constants must be symmetric: $\zeta_{ik} = \zeta_{ki}$. While cargo and streptavidin molecules exhibit Brownian motion and relax to a thermal equilibrium state with vanishing fluxes $\boldsymbol{j}_c = \boldsymbol{j}_a = 0$, the MinD protein patterns are kept in a non-equilibrium steady state maintained by off-equilibrium chemical reactions (ATPase activity). Because the fluxes of passive cargo and streptavidin molecules vanish, there is also no need to consider a coupling $\zeta_{ac}$ between them. Furthermore, Eq. S7 shows that the self-coupling coefficients $\zeta_{ii}$ are irrelevant for the mean field dynamics. For single molecules, however, such a self-coupling should lead to a density-dependent self-diffusion coefficient, as has been observed for MinD by Loose et al.[15] using single-particle tracking.

Since the fluxes of the passive particles, cargo and streptavidin molecules, vanish ($\boldsymbol{j}_c = \boldsymbol{j}_a = 0$), the fluxes of the MinD proteins are given by

$$\boldsymbol{j}_p = -\frac{c_p\,\nabla\mu_p}{\xi_p + c_c\,\zeta_{pc} + c_s\,\zeta_{ps}}, \qquad \text{(S8)}$$

where the chemical potential gradient of the MinD proteins is given by Eq. S2c. After inserting Eq. S8 back into the force balance equation, Eq. S7, one obtains the following relations between the externally maintained chemical potential gradients of the active particles (MinD proteins) and the induced chemical potential gradients of the passive particles (streptavidin and cargo molecules), respectively:

$$\nabla\mu_c = \zeta_{pc}\,\boldsymbol{j}_p = -\frac{c_p\,\zeta_{pc}}{\xi_p + c_c\,\zeta_{pc} + c_s\,\zeta_{ps}}\,\nabla\mu_p, \qquad \text{(S9a)}$$

$$\nabla\mu_s = \zeta_{ps}\,\boldsymbol{j}_p = -\frac{c_p\,\zeta_{ps}}{\xi_p + c_c\,\zeta_{pc} + c_s\,\zeta_{ps}}\,\nabla\mu_p. \qquad \text{(S9b)}$$

Here, the chemical potential gradient of the cargo molecules is given by Eq. S3c, the chemical potential gradient of streptavidin molecules is given by Eq. S2b, and the chemical potential gradient of the MinD proteins is given by Eq. S2c. Analogous to our numerical solution of the

Flory-Huggins model (see section 2 “Flory-Huggins theory of mixing: an equilibrium picture”), we formulated Eqs. S9a and S9b as a 1D boundary-value problem in a domain of size $L \equiv 1$, by introducing the two additional fields $\Theta_o = \frac{1}{L}\int_0^x dy\, \theta_o(y)$ and $\Theta_s = \frac{1}{L}\int_0^x dy\, \theta_s(y)$ and their respective boundary conditions $\Theta_s(0) = 0$, $\Theta_s(L) = \overline{\theta_s}(n)$ , $\Theta_o(0) = 0$, and $\Theta_o(L) = \overline{\theta_o}(n)$. As before, we imposed the (externally maintained) distribution of MinD proteins as defined by Eq. S6.

In the absence of mutual friction between the macromolecules ($\zeta_{ik} = 0$), Eq. S7 reduces to the Flory-Huggins model ($\boldsymbol{\nabla}\mu_c = \boldsymbol{\nabla}\mu_s = 0$) which implies weak cargo redistribution in a static gradient of active proteins. In contrast, in the presence of (frictional) coupling ($\zeta_{ik} \neq 0$) between cargo molecules and MinD protein fluxes ($\boldsymbol{j}_p \neq 0$), cargo molecules are not only redistributed due to entropic unmixing effects, but in addition transported along protein gradients by these nonequilibrium protein fluxes. As consequence of this additional bias, cargo redistribution is significantly stronger than by equilibrium thermodynamic forces alone, which quantitatively explains our experimental data (Fig. 2b,c in the main text). We expect that the coupling constant $\zeta_{pc}$ between MinD and a specific cargo has a small contribution from the origami scaffold as well as from each of its $n$ streptavidin:

$$\zeta_{pc} = \zeta_{po} + n\, \zeta_{ps}. \tag{S10}$$

This implies that cargo transport should increase with the number of streptavidin integrated into the cargo, as observed in our experiments (Fig. 1d in the main text). We also expect that individual streptavidin molecules experience a significant coupling $\zeta_{ps}$ to MinD fluxes. Therefore, Maxwell-Stefan diffusiophoresis is not limited to the transport of large cargo but can also explain the transport of small molecules (with similar size as MinD proteins) as reported in this study (Fig. 1c) and previous ones[16,17].

##### 4. Analytic solution and fitting of reduced model

To further elucidate the mechanism underlying MinDE-induced transport, we simplified our theoretical model. Specifically, we neglected membrane saturation effects (second term in Eqs. S2a-c and S3b), so that the chemical potential of a particle with size $a_i$ reduces to $\mu_i \approx k_B T \ln(a_i c_i)$. Then, the effective force-balance equation, Eq. S7, takes the form of a generalized Fick’s law for the protein fluxes in the non-equilibrium steady state with a density-dependent diffusion coefficient $D_p(c_c, c_s)$:

$$\boldsymbol{j}_p \approx -\frac{k_B T}{\xi_p + c_c\,\zeta_{pc} + c_s\,\zeta_{ps}}\,\boldsymbol{\nabla} c_p \equiv -D_p(c_c, c_s)\,\boldsymbol{\nabla} c_p. \qquad \begin{pmatrix} \text{S11,} \\ \text{2 in main text} \end{pmatrix}$$

Because the number of free streptavidin is typically small (see section 1 "Cargo density is limited by abundance of streptavidin and origami"), we assumed that free streptavidin molecules do not significantly contribute to the dynamics, $c_s\,\zeta_{ps} \ll c_c\,\zeta_{pc}$. After inserting Eq. S11 back into the force balance equation, Eq. S7, one obtains the following relation between cargo molecule and MinD protein gradients:

$$\boldsymbol{\nabla} c_c = -\frac{c_c}{c_\times + c_c}\boldsymbol{\nabla} c_p, \qquad \text{(S12)}$$

where we defined the *interaction density* $c_\times = \xi_p/\zeta_{pc}$. In the equilibrium state, where the fluxes of cargo molecules vanish $\boldsymbol{j}_c = 0$, the distribution of the cargo molecules will be determined by the distribution of proteins, the abundance of molecules in the assay, and the interaction density $c_\times$. Therefore, the local cargo density will have the form $c_c(c_p, c_\times, \bar{c_c}, \bar{c_p})$. Thus, by using the chain rule of differentiation, one can fully eliminate all gradients from Eq. S12 to obtain the following ordinary differential equation:

$$\frac{\partial c_c}{\partial c_p} = -\frac{c_c}{c_\times + c_c}, \qquad \text{(S13)}$$

Alternatively, one can also obtain the ordinary differential equation S13 by integrating Eq. S12 over an arbitrary infinitesimal line segment $\boldsymbol{n}\,ds$, and perform a change of variables $ds\,\boldsymbol{n}\cdot\boldsymbol{\nabla} c = dc$. Eq. S13 can be solved by integration, and yields the following relationship between the cargo molecule density and the MinD protein density:

$$c_c(c_p) = c_\times\,W_0\left[\frac{c_c(0)}{c_\times}\exp\left(\frac{c_c(0) - c_p}{c_\times}\right)\right]. \qquad \begin{pmatrix} \text{S14,} \\ \text{3 in main text} \end{pmatrix}$$

Here, $W_0$ refers to the principal branch of the Lambert W-function, which is defined as the inverse function of $f(x) = x\,e^x$ (cf. illustration "Lambert W-function"). In terms of fluorescence intensities, $I_c = \alpha_c c_c$ and $I_p = \alpha_p c_p$, Eq. S14 can be rewritten as

$$I_c(I_p) = r\,I_\times\,W_0\left[\frac{I_c(0)}{r\,I_\times}\exp\left(\frac{I_c(0)}{r\,I_\times} - \frac{I_p}{I_\times}\right)\right], \qquad \text{(S15)}$$

where we have defined the fluorescence ratio $r = \alpha_c/\alpha_p$ and the typical MinD intensity corresponding to the interaction density $I_\times = \alpha_p c_\times$. In our experiments, we controlled the abundance of all fluorescent molecules. Thus, by measuring the average fluorescence intensity

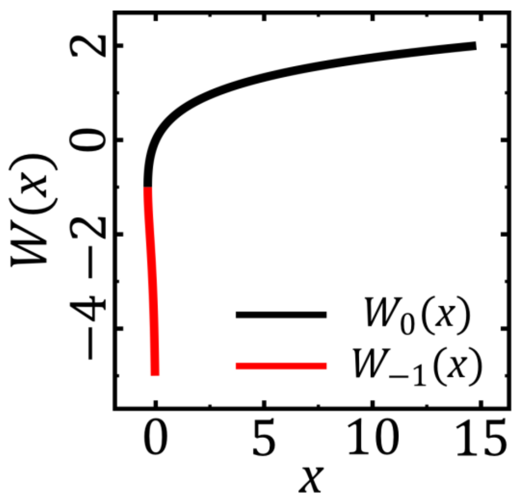

of the DNA origami (7 000, in arbitrary units) and knowing their average density (313 µm$^{-2}$ in assuming that all DNA origami bind to the membrane), we estimated the fluorescence yield of a DNA origami/cargo molecule as $\alpha_c \approx 24$ µm$^2$. The MinD density in the MinD maxima reaches a value of about 13 200 µm$^{-2}$ [15,16]. Thus, by measuring the fluorescence intensity in the MinD maxima (20 000, in arbitrary units), we estimated the fluorescence yield of a MinD protein as $\alpha_p \approx 1.5$ µm$^2$. This results in the following fluorescence ratio: $r \approx 15$.

#### 5. Diffusion coefficient of cargo molecules

To further validate our model with corresponding experiments, we aimed to determine the diffusion coefficient of cargo molecules. Analogously to section 4 "Analytic solution and fitting of reduced model", we neglected membrane saturation effects, so that the chemical potential of a particle with size $a_i$ reduces to $\mu_i \approx k_B T \ln(a_i c_i)$. Furthermore, because the number of free streptavidin is typically small (see section 1 "Cargo density is limited by abundance of streptavidin and origami"), we neglected the density of free streptavidin molecules ($c_s \approx 0$). Then, the effective force-balance equation, Eq. S7, takes the form of a generalized Fick's law for the cargo fluxes, where gradients in the cargo molecule density ($\boldsymbol{\nabla} c_c$) relax with a density-dependent diffusion coefficient $D_c(c_p)$:

$$\boldsymbol{j}_c \approx -\frac{k_B T}{\xi_c + c_p\,\zeta_{pc}}\,\boldsymbol{\nabla} c_c \equiv -D_c(c_p)\,\boldsymbol{\nabla} c_c. \qquad \text{(S16)}$$

By substituting the typical interaction density $c_\times = \xi_p/\zeta_{pc}$, the diffusion constant of cargo molecules in the dilute limit $D_c^0 = k_B T/\zeta_c$, and the diffusion constant of MinD proteins in the dilute limit $D_p^0 = k_B T/\zeta_p$, one obtains:

$$D_c(c_p) \approx \frac{D_c^0}{1 + (D_c^0/D_p^0)(c_p/c_\times)} \,. \tag{S17}$$

We measured the diffusion constant of cargo-2 and of cargo-42 in the dilute limit. From measurements performed by Loose et al.[15], we estimated the diffusion constant of membrane-bound MinD proteins in the dilute limit as $D_p^0 \approx 0.425\ \mu m^2 s^{-1}$.

##### 6. Simulations with multiple cargo species

Simulations with multiple cargo species were performed analogously to simulations with one cargo species. Specifically, we added new species to the expressions for the chemical potentials (Eqs. S2a-c, Eq. S3b; additional chemical potential for the new species), to the Flory-Huggins model (Eqs. S4a,b; additional PDE for the new species), and to the Maxwell-Stefan model (Eqs. S9a,b; additional PDE for the new species). Furthermore, we assumed equal abundance of all different cargo species on the membrane.

#### II. Discussion of alternative thermodynamic transport mechanisms

##### 1. Depletion forces cannot explain cargo transport

Depletion forces arise from a classical entropic effect, where finite-sized molecules (like MinD) can access a larger spatial region if larger molecules (like DNA-Origami) 'clump' together[18]; see Fig. 5 for an illustration. This results in effective (Asakura-Oosawa)[18] depletion forces that act on the larger molecules and which are proportional to the concentration of the smaller molecules, $c_p$. Phenomenologically, one can represent these depletion forces as a *negative pressure*, $p \propto -c_p$. If the concentration of the smaller molecule, $m_p$, is spatially heterogeneous, then this will result in effective pressure gradients, $-\boldsymbol{\nabla} p \propto \boldsymbol{\nabla} c_p$. Consequently, one would expect that depletion forces lead to an accumulation of cargo molecules in MinD-rich regions. Because this expectation contradicted our experiments, we concluded that depletion forces play no significant role for cargo transport. Furthermore, we did not observe depletion-force-induced aggregation of cargo molecules when MinD was homogenously distributed (Fig. S3).

##### 2. Static volume exclusion cannot explain cargo transport

Suppose that membrane-bound MinD proteins act as static obstacles of size $a_p$ and surface density $c_p$. Then, such obstacles locally occupy a fraction $\theta_p = a_p c_p$ of the surface, thereby reducing space accessible by the cargo molecules. In thermal equilibrium, cargo molecules spread uniformly across the accessible space, which implies $c_c \propto \theta_{\text{free}} = (1 - \theta_p)$ for the cargo molecule density. Formally, this can be seen by solving for the steady-state solution of cargo diffusing in a porous medium:

$$\partial_t c_c = \nabla[D_c \, \theta_{\text{free}} \, \nabla(c_c/\theta_{\text{free}})] \, . \tag{S18}$$

However, this implies that the resulting distribution of cargo molecules does not depend on any intrinsic features of the cargo molecules, contradicting our experimental observations (Fig. 1c-f). We conclude that MinD proteins do not act as static obstacles for the cargo molecules.

##### 3. Significant impact on reaction kinetics is unlikely

As the MinDE distribution was influenced by the presence of cargo, we wondered whether cargo may change the kinetic (un)binding rates of MinDE. To answer this question, we analyzed the average fluorescence intensity of the patterns, i.e. the membrane density of the molecules. While we found that the density of streptavidin and MinD were relatively similar for all conditions, the density of membrane-bound DNA origami decreased by roughly 44% when we increased the number of streptavidin building blocks from 1 to 42 (Fig. S5b-d). The latter suggested that the average density of membrane-bound streptavidin, which remained unaffected, is the limiting factor for binding of DNA origami to the membrane (see SI I. 1. for details). As the presence of cargo did not change the average membrane density of MinD (Fig. S5b), it is unlikely to significantly affect their (un)binding rates. Furthermore, cargo always accumulated in regions where both the MinD density and thus protein recruitment to the membrane are already low, and is thus unlikely to significantly hinder protein (un)binding.

Cargo and streptavidin molecules have a strong membrane affinity and negligible detachment rates. Therefore, MinD-induced detachment of cargo or streptavidin from the membrane is highly unlikely.

### Supplementary Figures

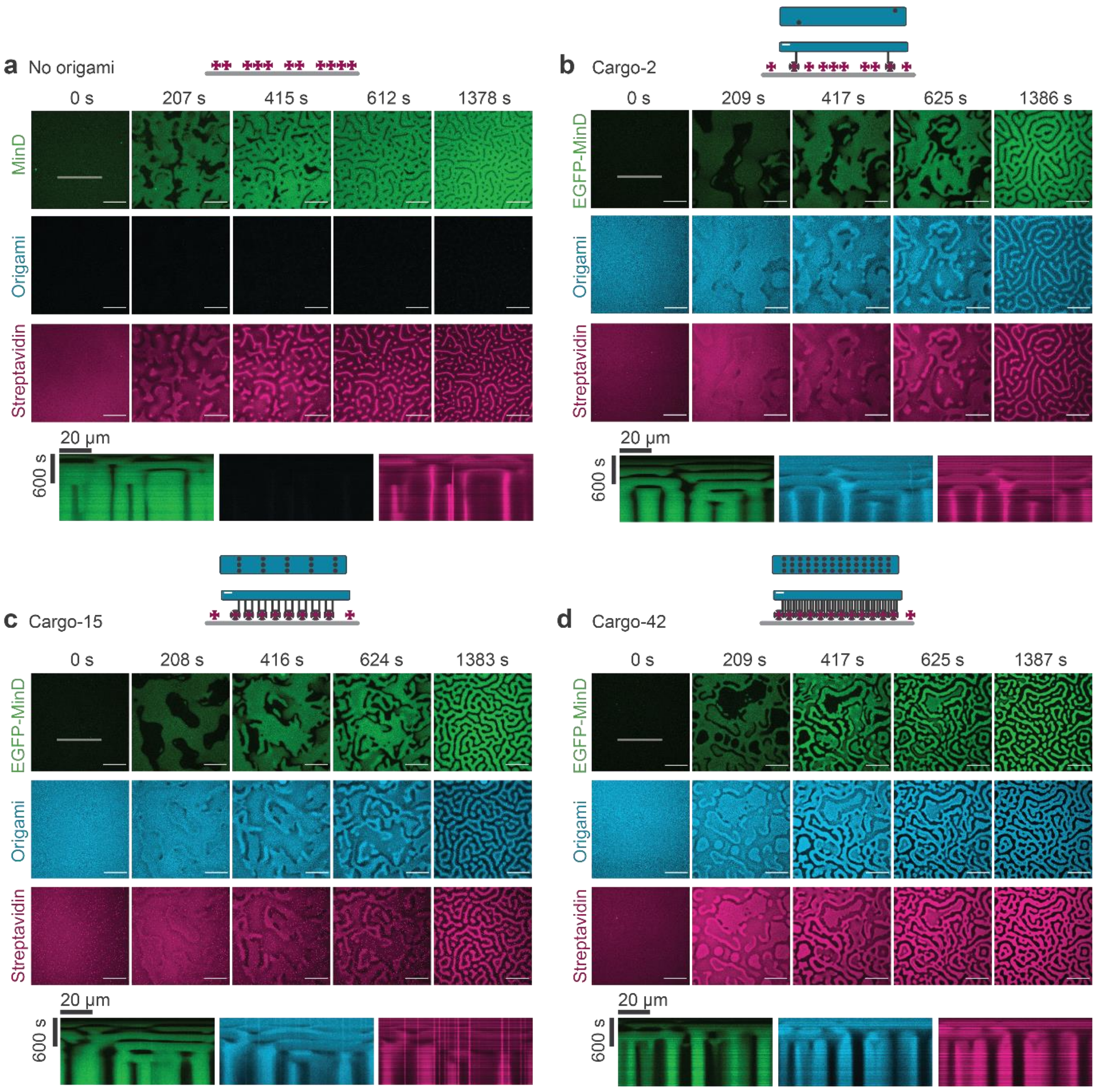

**Supplementary Figure S1: MinDE induces pattern formation of cargo from an initially homogenous state**. Representative time-series and kymograph (along line selection) of MinDE self-organization inducing patterns of DNA origami and streptavidin when **a**, no origami, **b**, cargo-2, **c**, cargo-15 and **d**, cargo-42 is present (1 µM MinD (30% EGFP-MinD), 1.5 µM MinE-His, in absence or presence of 0.1 nM origami-Cy5 with 2, 15 or 42 biotinylated oligonucleotides, Alexa568-streptavidin). Note, that panel b is identical to Figure 1b. Scale bars: 50 µm

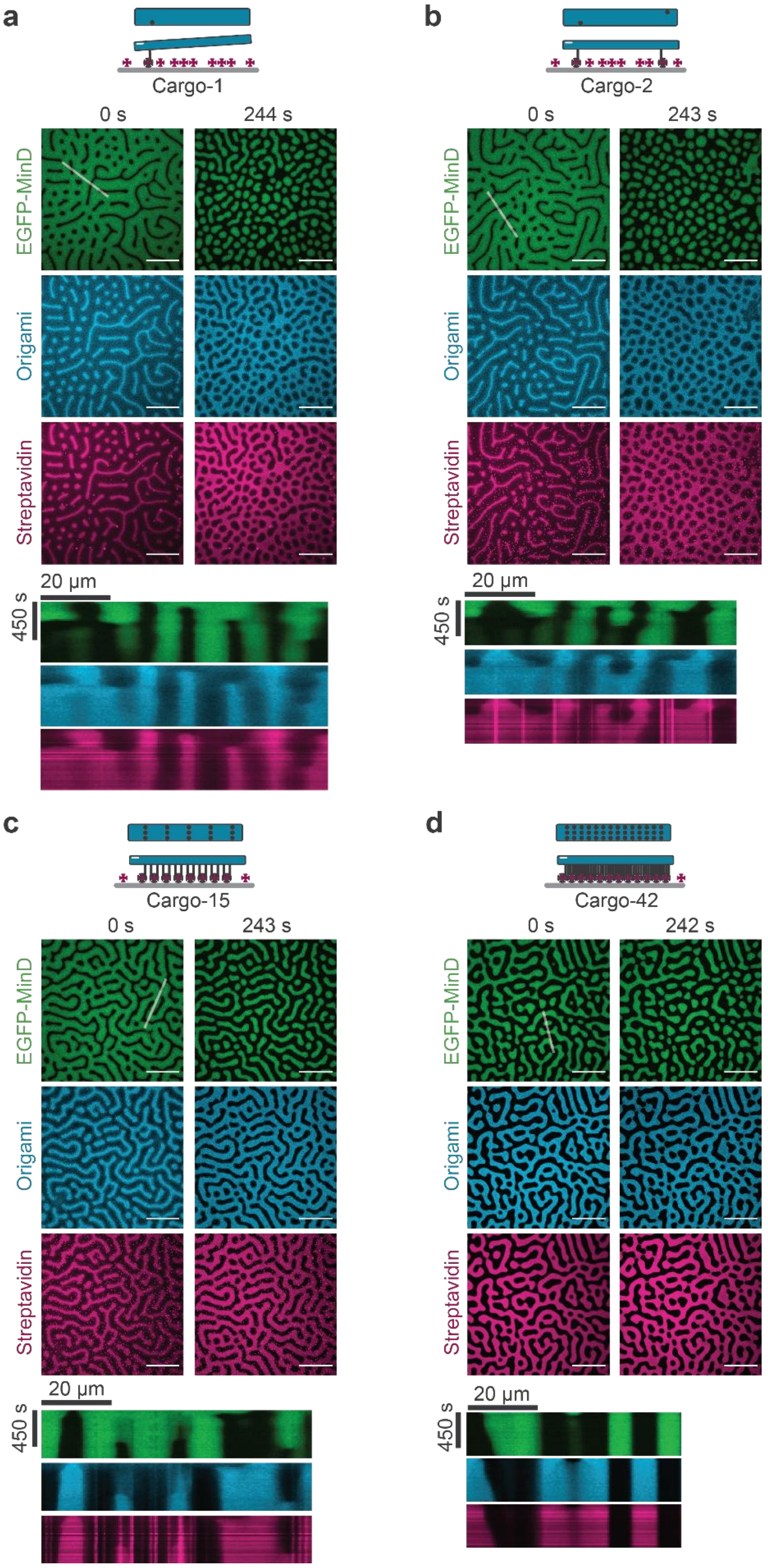

**Supplementary Figure S2: Changes in the MinDE patterns are mimicked by cargo**. Representative time-series and kymograph showing changes to MinD and cargo molecule patterns in presence of **a**, cargo-1, **b**, cargo-2, **c**, cargo-15 and **d**, cargo-42 (1 µM MinD (30% EGFP-MinD), 1.5 µM MinE-His, origami-Cy5 with 1, 2, 15 or 42 biotinylated oligonucleotides, Alexa568- streptavidin) upon addition of more MinE (addition of 1.5 µM MinE-His). MinE addition directly before t = 0 s. Scale bars: 50 µm

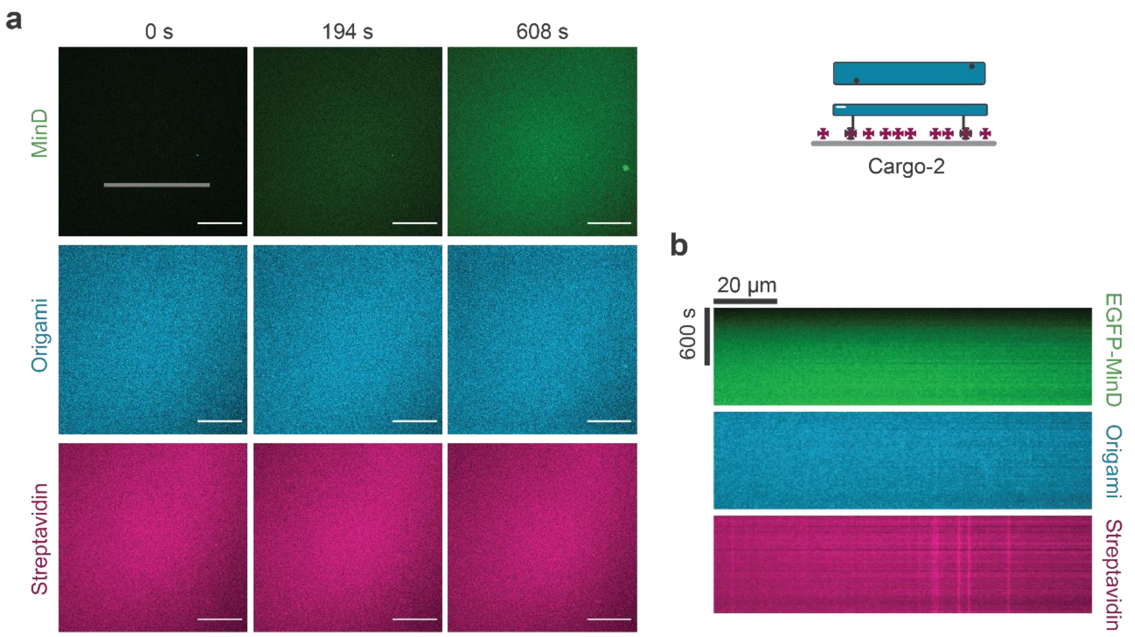

**Supplementary Figure S3: Pattern formation of DNA origami and streptavidin depends on active self-organization by MinDE. a**, Representative time-series of MinD membrane binding on SLBs in the presence of cargo-2 and ATP, but in the absence of MinE (1 µM MinD (30% EGFP-MinD), 0.1 nM origami-Cy5 with 2 biotinylated oligonucleotides, Alexa568-streptavidin, ATP). Scale bars: 50 µm. **b**, Kymographs of the line selection indicated in a.

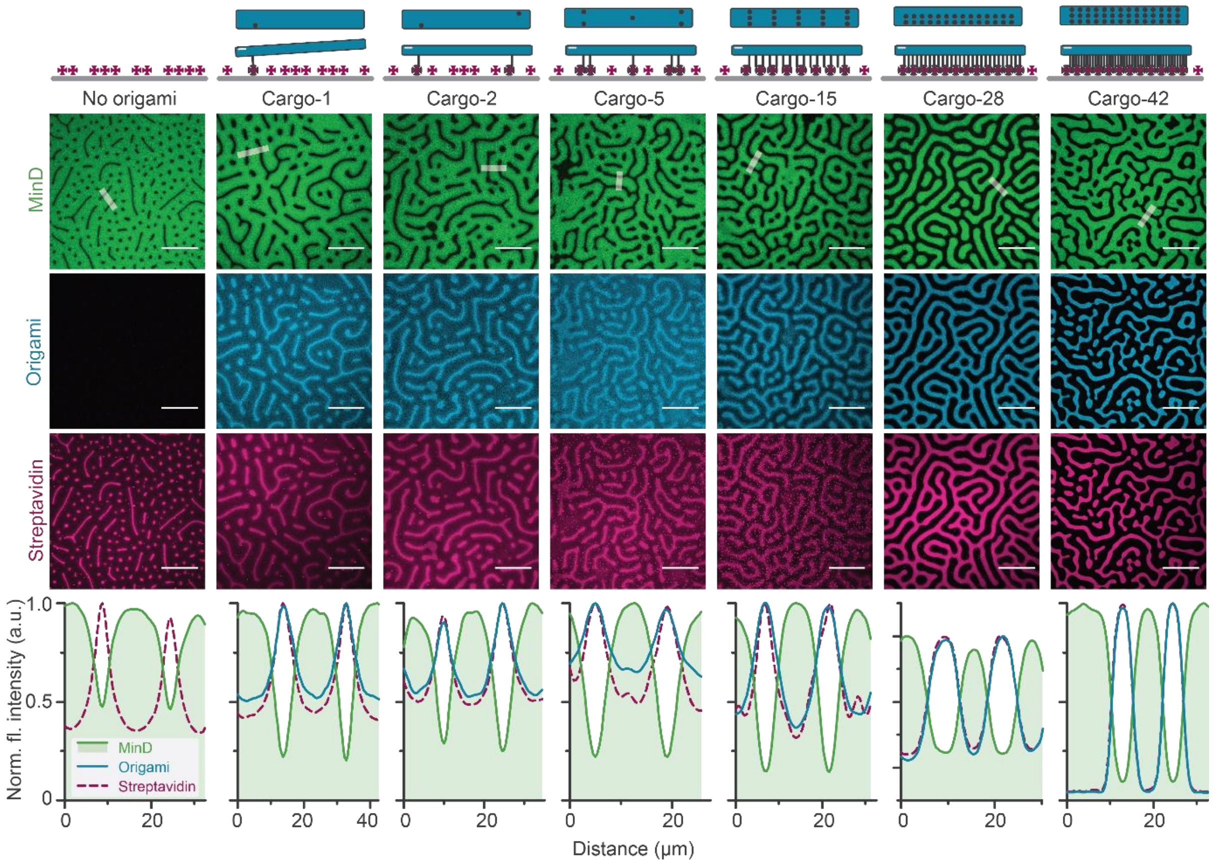

**Supplementary Figure S4: Extent of the MinDE-driven cargo unmixing depends on the effective size of the cargo**. Representative images and fluorescence intensity line plots (smoothed) of established MinDE labyrinth patterns and anti-correlated DNA origami and streptavidin patterns when no origami or cargo-1, cargo-2, cargo-5, cargo-15, cargo-28, cargo-42 is present (1 µM MinD (30% EGFP-MinD), 1.5 µM MinE-His, with or without 0.1 nM origami-Cy5 with *n* biotinylated oligonucleotides, Alexa568-streptavidin). Scale bars: 50 µm

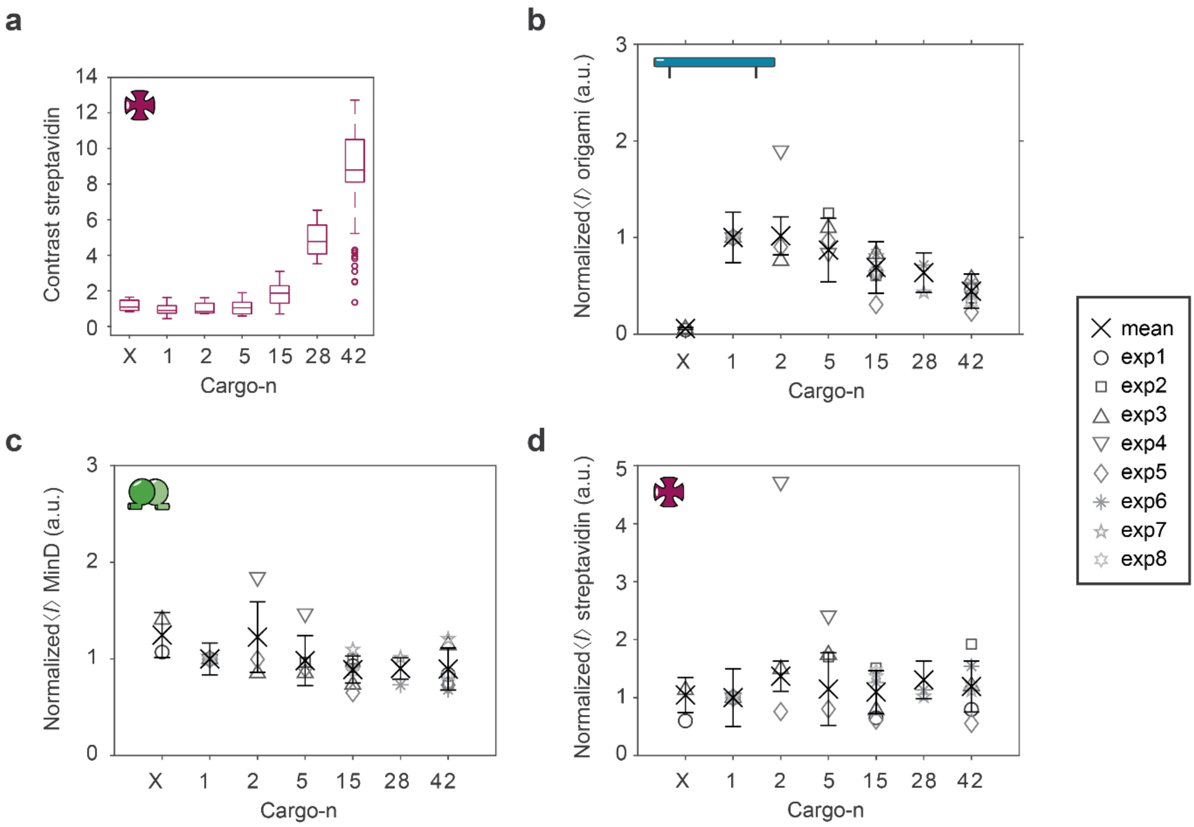

**Supplementary Figure S5: Contrast of patterns increases while membrane densities remain the same or decrease with increasing numbers of streptavidin a**, Contrast of the resulting streptavidin patterns increases with increasing number of streptavidin. Box plot lines are median, box limits are quartiles 1 and 3, whiskers are 1.5× interquartile range (IQR) and points are outliers. Mean fluorescence intensity of **b**, DNA origami, **c**, EGFP-MinD and **d**, Alexa568-streptavidin and of the full image, normalized to the intensity of experiments containing cargo-1, of patterns formed when no origami or cargo-1, cargo-2, …, cargo-42 is present. Cross and error bars represent the mean value and standard deviation of two or more independent experiments with in total N images per condition N(No origami)=32, N(Cargo-1)=112, N(Cargo-2)=41, N(Cargo-5)=48, N(Cargo-15)=110, N(Cargo-28)=32, N(Cargo-42)=98. Scale bars: 50 µm

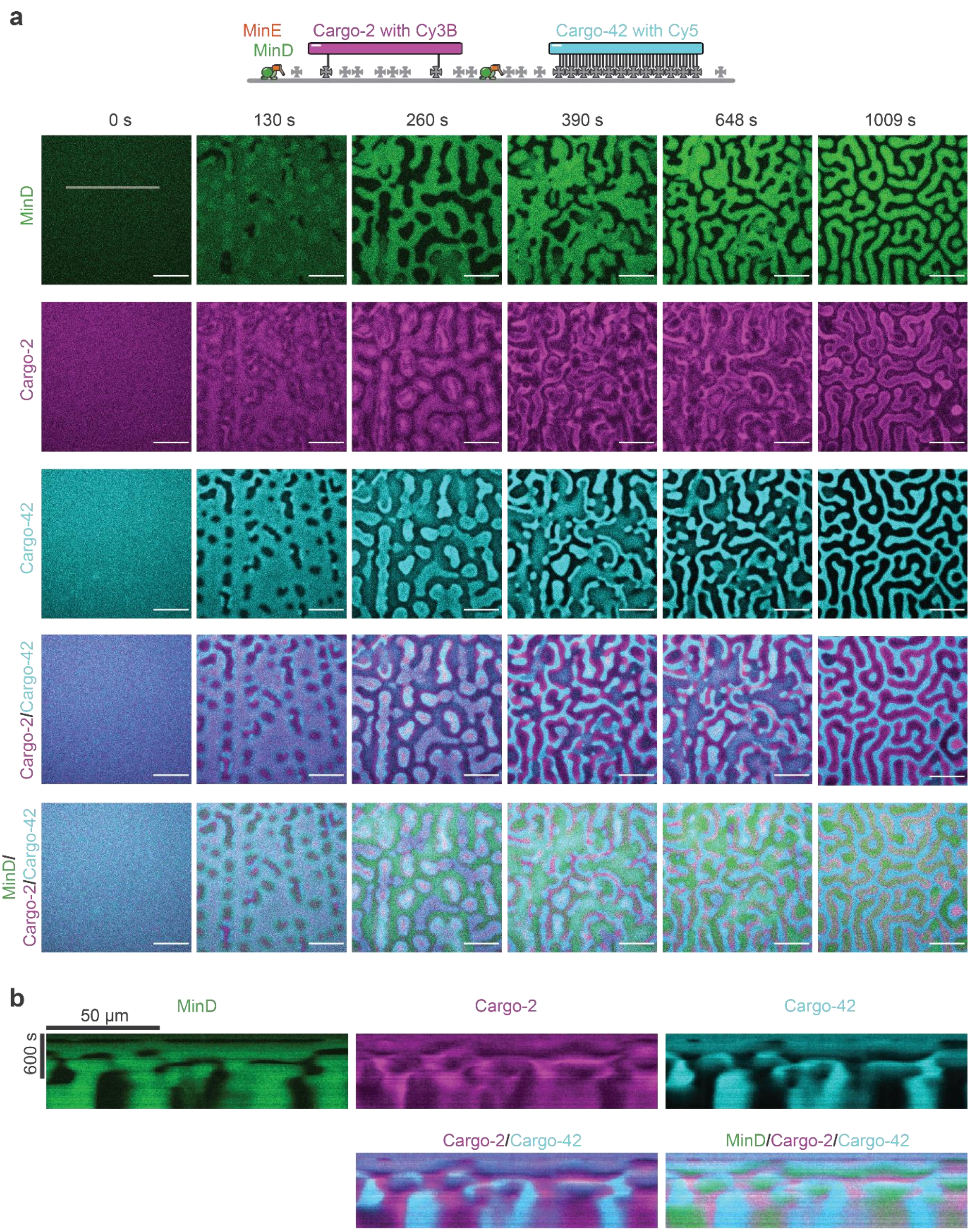

**Supplementary Figure S6: MinDE induces pattern formation of two distinct cargoes, cargo-2 and cargo-42, from an initially homogenous state**. **a**, Representative time-series of MinD, cargo-2 and cargo-42 pattern formation. ATP is added to start self-organization directly before t=0 s (1 µM MinD (30% EGFP-MinD), 1.5 µM MinE-His, 50 pM origami-Cy3B with 2 biotinylated oligonucleotides, 50 pM origami-Cy5 with 42 biotinylated oligonucleotides, non-labelled streptavidin). Scale bars: 50 µm **b**, Kymographs of the line selection shown in a.

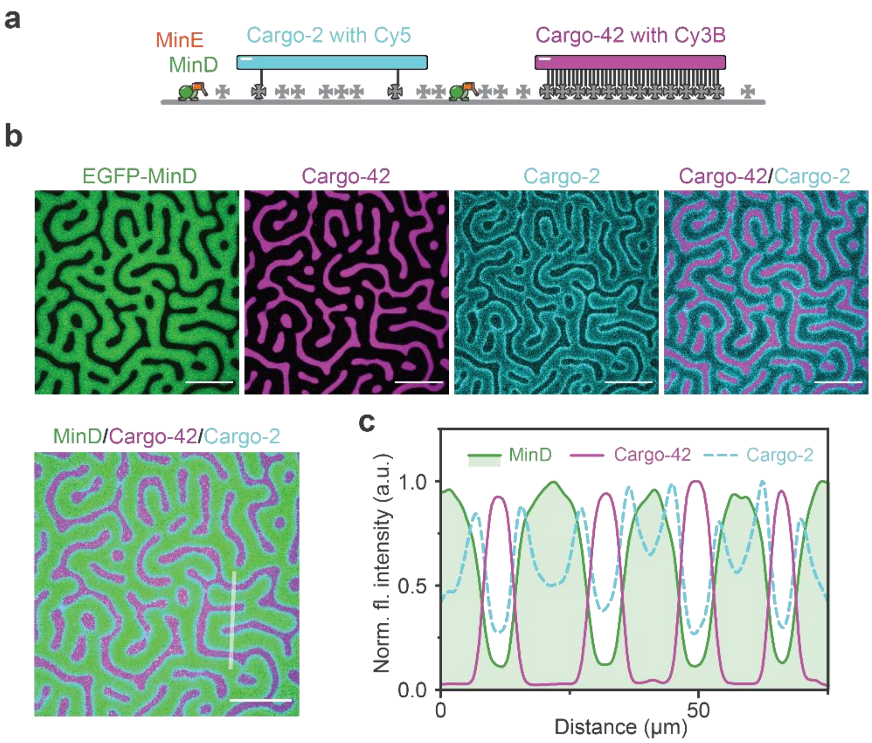

**Supplementary Figure S7: Spatial sorting of cargo-2 and cargo-42 by MinDE also occurs when dyes are swapped**. **a**, Schematic of the experimental setup. MinDE self-organization was performed in presence of two different cargo species with distinct fluorescent labels, cargo-2 with Cy5 and cargo-42 with Cy3B. **b**, Representative images and **c**, lineplots of MinDE-induced sorting of cargo-2 and cargo-42 (1 µM MinD, 1.5 µM MinE-His, 50 pM origami-Cy5 with 2 biotinylated oligonucleotides, 50 pM origami-Cy3B with 42 biotinylated oligonucleotides, non-labelled streptavidin). Experiment was performed three times under identical conditions. Scale bars: 50 µm

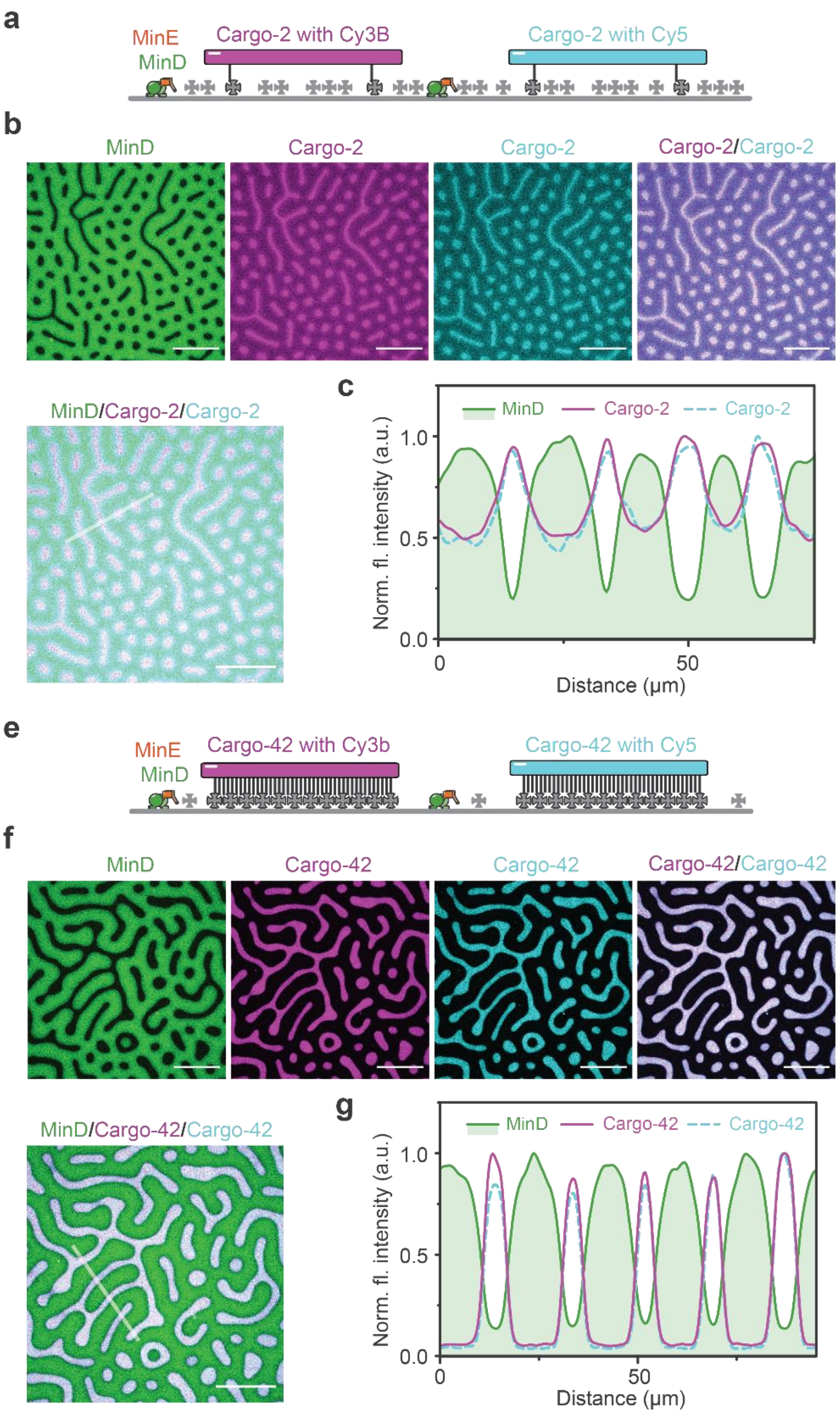

**Supplementary Figure S8: Spatial sorting is not caused by the labelling of cargo.** MinDE-induced distributions of differentially labelled, but otherwise identical cargo are superimposable. **a**, and **f**, Schematic of the experimental setup: two identical cargoes are labelled with distinct dyes. Pattern formation is induced by addition of MinDE (1 µM MinD, 1.5 µM MinE-His, 50 pM origami-Cy5 and 50 pM origami-Cy3B, non-labelled streptavidin). Representative images and line plot of pattern formation for presence of **b**, **c**, two differently labelled cargo-2 and **e**, **f**, two differently labelled cargo-42. Experiment was performed two times under identical conditions. Scale bars: 50 µm

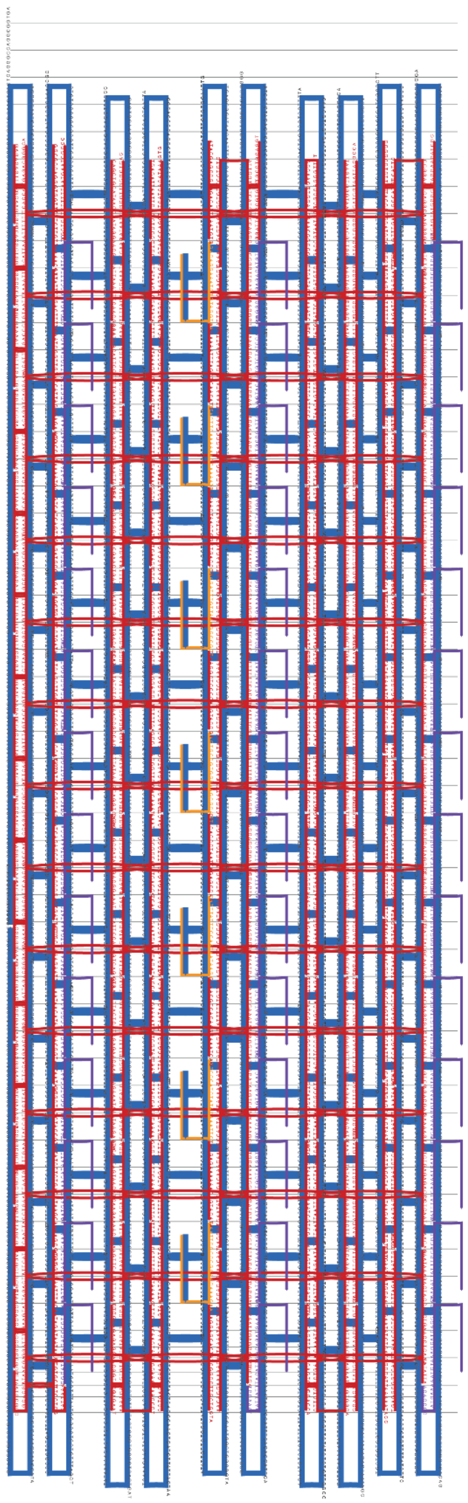

**Supplementary Figure S9. Design of the elongated 20-helix bundle DNA origami with 42 handle positions.** The dye-modified and connector oligonucleotides required for fluorescence detection are highlighted in orange, the 42 possible positions for incorporation of biotinylated oligonucleotide handles for binding to streptavidin in purple, core staples in black and the M13 p7249 scaffold is coloured in blue.

### Supplementary Table

**Supplementary Table 1: Model Parameters and variables.** Overview of the parameters and all dependent variables used in Flory-Huggins type and Maxwell-Stefan type theories.

| *Parameter* | **Value** | **Explanation** |
|---|---|---|
| $\theta_{\times}^{0}$ | 10 | Interaction parameter between origami scaffold and MinD, in terms of MinD coverage |
| $\theta_{\times}^{s}$ | 2.5 | Interaction parameter between streptavidin and MinD, in terms of MinD coverage |
| $n$ | 2—42 | Number of streptavidin blocks attached to origami |
| ***Estimated from experiments:*** | | |
| $\overline{\theta_p}$ | 0.20 | Average MinD surface coverage. Derived from measured densities[15,16]. |
| $\overline{\theta_{o+}}$ | 0.56 | Average origami surface coverage (distal), if all available DNA origami in the assay (0.1 nM) were to bind to the available streptavidin molecules. |
| $\overline{\theta_{s+}}$ | 0.15 | Average surface coverage of available streptavidin molecules (free + bound). Derived from measured densities[16]. |
| ***Dependent variables:*** | | |
| $\overline{\theta_o}$ | $\min(\overline{\theta_{o+}}, \overline{\theta_{s+}}\, a_o/a_c)$ | Average origami surface coverage (distal), which is limited by the density of available streptavidin molecules. |
| $\overline{\theta_s}$ | $\overline{\theta_{s+}} - \overline{\theta_o}\, a_c/a_o$ | Average surface coverage of free streptavidin molecules. |

### Supplementary Note

**Supplementary Note 1:**

The DNA origami rod is located about 5-11 nm above the membrane, as it is bound to the membrane via several spacers: the dsDNA oligonucleotide linker, a TEG-biotin moiety and the streptavidin molecule. The length of the dsDNA linker can be estimated to about 6.1 nm taking into account the rise per basepair (bp) of 0.34 nm of B-DNA[19]. The oligonucleotide is connected to the anchoring biotin moiety via TEG with a length of about 1.4 nm[20]. And thus the total linker length can be estimated to be about 7 nm. As the persistence length of dsDNA is about 50 nm[19,21,22] the dsDNA linker is rigid. However, the connection to the DNA origami is only single stranded giving the dsDNA linkers freedom for bending[23]. The height of membrane-bound streptavidin was measured to be ~4 nm[24] in good agreement with measurements from EM/crystal structures[25,26].
